# Oral glucose feeding enhances adherence of quiescent lymphocytes to fibronectin via non-canonical insulin signalling

**DOI:** 10.1101/2021.10.13.464163

**Authors:** Abhiram Charan Tej Mallu, Sivapriya Sivagurunathan, Debasish Paul, Hobby Aggarwal, Abel Arul Nathan, Mahalakshmi M. Ravi, Ramanamurthy Boppana, Kumaravelu Jagavelu, Manas Kumar Santra, Madhulika Dixit

## Abstract

Impaired glucose metabolism is associated with chronic inflammation, aberrant immunity and anomalous leukocyte trafficking. Conversely, infusion of functional immune cells restores glucose metabolism. Despite being exposed to periodic alterations in blood insulin levels upon fasting and feeding, studies exploring the physiological effects of these hormonal changes on quiescent circulating lymphocytes are missing. Here we find that oral glucose load in healthy men and mice enhance adherence of circulating peripheral blood mononuclear cells (PBMCs) to fibronectin. This led to increased homing of post-load PBMCs to injured blood vessels. Cell culture based experiments on Jurkat-T cells and PBMCs demonstrated that insulin elicits these adhesive effects through a non-canonical signalling involving insulin growth factor-1 receptor (IGF-1R) and phospholipase C gamma-1 (PLCγ-1) mediated activation of integrin β1. Our findings point to the relevance of post-prandial insulin spikes in regulating homing of circulating T-cells to various organs for tissue repair and immunity.

**Graphical Abstract:** Insulin mediates fibronectin adherence of lymphocytes through non-canonical signalling. Insulin mediates auto-phosphorylation of IGF-1 receptor at Tyr^1135^ leading to activation of PLC-γ1 through Tyr783 phosphorylation, which in turn leads to the activation of integrin β1 through intracellular calcium to ultimately enhance adhesion of quiescent lymphocytes to fibronectin.

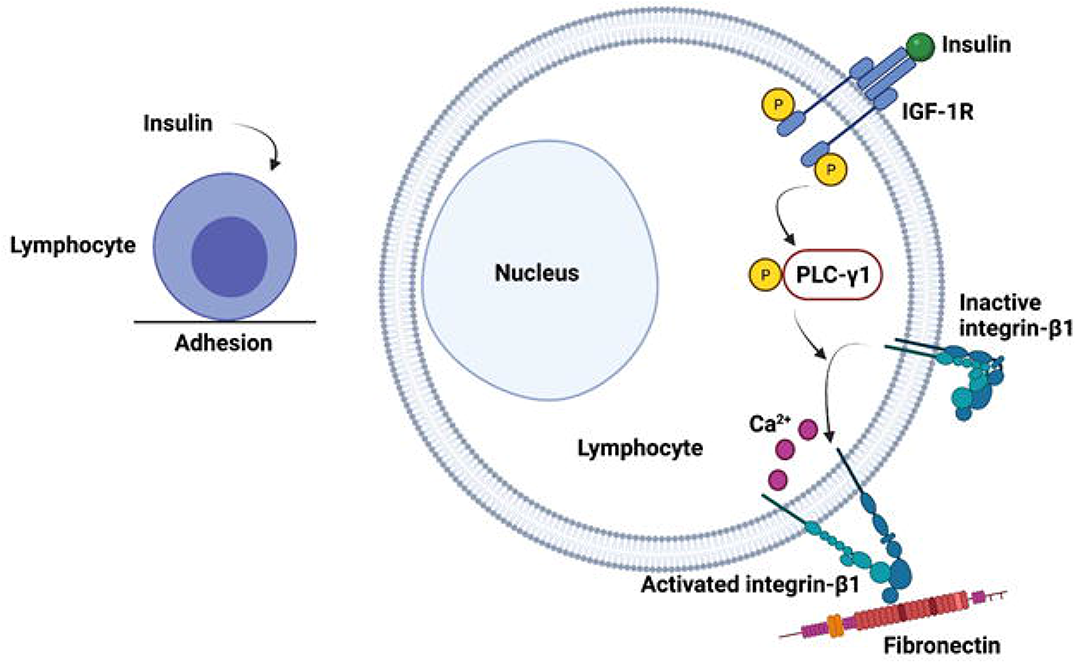

## 1. Introduction

Metabolism and immunity exhibit coevolution and intricate cross-talk in order to regulate each other. For instance, activation of either neutrophils or T-lymphocytes upon antigen presentation, is accompanied by increased glycolysis and insulin signaling [1, 2]. Additionally, development of immunity in the new born through mother’s breast milk, epigenetic modifications during T-cell differentiation, or gross reprogramming of gene expression in immune cells during insulin resistance, all support this inter-dependence [3–9]. It is thus not surprising to note that impaired glucose metabolism in diabetes and obesity is often associated with chronic low-grade inflammation, altered cytokine profiles, aberrant immune response and sub-optimal trafficking of leukocytes and stem cells [10–13]. These aberrations in turn predispose individuals to infections, atherosclerosis and cancer [14, 15]. In contrast, adoptive transfer of functional Treg cells restores glucose tolerance and mitigates insulin resistance in high fat diet induced obesity [16, 17]. Although the aforementioned studies demonstrate that chronic metabolic impairments disrupt immunity, and conversely, infusion of functional immune cells restore systemic glucose metabolism; studies exploring the physiological influence of acute and daily changes in blood glucose and insulin on the function of circulating immune cells are scanty.

Calorie restrictions in the form of fasting or fasting mimicking diets, attenuate immune senescence and inflammation, reverse myeloid to lymphoid cell ratio and promote tissue repair [3, 18]. These diets decrease organ weights with refeeding aiding the repopulation of cells in respective vascular beds [18]. The composition of diet along with frequency and timing of consumption is reported to dictate immune regulation [19–21]. For instance, high protein diet promotes expression of genes involved in immune activation, antigen presentation and cytokine signaling, whereas, saturated fatty acids elicit innate immune response [22, 23]. In healthy individuals, acute dietary challenges in the form of oral glucose load following over-night fasting, triggers a well-controlled inflammatory response. These studies are however restricted to observing changes either in serum biochemistry or in gene expression of peripheral blood mononuclear cells (PBMCs) [8,24–26]. Intriguingly, studies addressing the effects of oral glucose load on leukocyte function and trafficking remain elusive. Despite being periodically exposed to altering insulin levels upon fasting and feeding, it is unclear if naïve circulating T-lymphocytes, which do not express insulin receptors on their surface [27], respond to post-prandial changes in insulin spikes. Hence, the primary objective of this study was to determine the effects of oral glucose load and insulin signaling on circulating PBMCs and Jurkat T-lymphocytes.

## 2. Results

### 2.1. Clinical characteristics of the study participants

Twenty drug naïve healthy Asian Indian men, 20-45 years of age, with no prior history of diabetes, were screened through a standard 75g oral glucose tolerance test (OGTT). Following application of inclusion and exclusion criteria as listed in the methods section, a total of sixteen healthy normal glucose tolerant (NGT) subjects were recruited in the study. Anthropometric and fasting biochemical data of the study subjects are summarized in table 1. Individuals were fairly young with a mean age of 29.2 years. They had normal blood pressure and a mean BMI of 26. Fasting blood glucose and insulin levels were in the normal range. The average HOMA-IR value for the recruited subjects was 1.8 indicating good insulin sensitivity and none of the subjects exhibited dyslipidemia (data not shown). It is to be noted that none of the recruited subjects were on any medication at the time of sample collection.

**Table 1:** Anthropometric and biochemical parameters of study participants. Data is represented as mean ± S.E.M.

| **Parameters** | **NGT (N=16)** |
| --- | --- |
| Age | 29.2 ± 1.15 |
| BMI (Kg/m<sup>2</sup>) | 26.0 ± 0.91 |
| Systolic BP (mm Hg) | 123.3 ± 2.05 |
| Diastolic BP (mm Hg) | 76.5 ± 1.57 |
| Fasting Plasma Glucose (mg/dl) | 87.1 ± 1.92 |
| Fasting Serum Insulin (μIU/ml) | 8.5 ± 0.80 |
| HOMA-IR | 1.8 ± 0.19 |

*BMI: body mass index, BP: blood pressure, HOMA-IR: Homeostatic model assessment of insulin resistance.*

### 2.2. Biochemical parameters following water and oral glucose load

In order to understand the specific effects of glucose load on PBMC function, prior to the 75g oral glucose tolerance test (OGTT) day, fasting and 2-hour post water load samples from the same study subjects were collected to serve as control. Hence, ten NGT subjects were called again on two different days to collect the blood samples at fasting and at 2-hours following intake of either 300ml water or 75gm glucose suspended in water. A gap of minimum 3 days was followed between the water and the glucose load days. All subjects underwent over-night fasting of 8-12 hours prior to collection of fasting samples on all days. Neither the plasma glucose levels, nor the serum insulin levels changed upon water consumption in individuals post water load (Fig. 1A and 1B). For the OGTT day, the 1-hour post load glucose levels were significantly higher compared to fasting, while the 2-hour post load glucose levels returned back to the normal range, indicating good blood-glucose clearance in study subjects (Fig. 1A). The fasting insulin was in the normal range and as expected, the 1- and 2-hour post load insulin levels were significantly higher compared to fasting in response to oral glucose feeding (Fig. 1B). The total and differential leukocyte count at fasting and at 2-hour post water and glucose load did not vary across conditions (Supplementary figure 1A).

**Figure 1:**
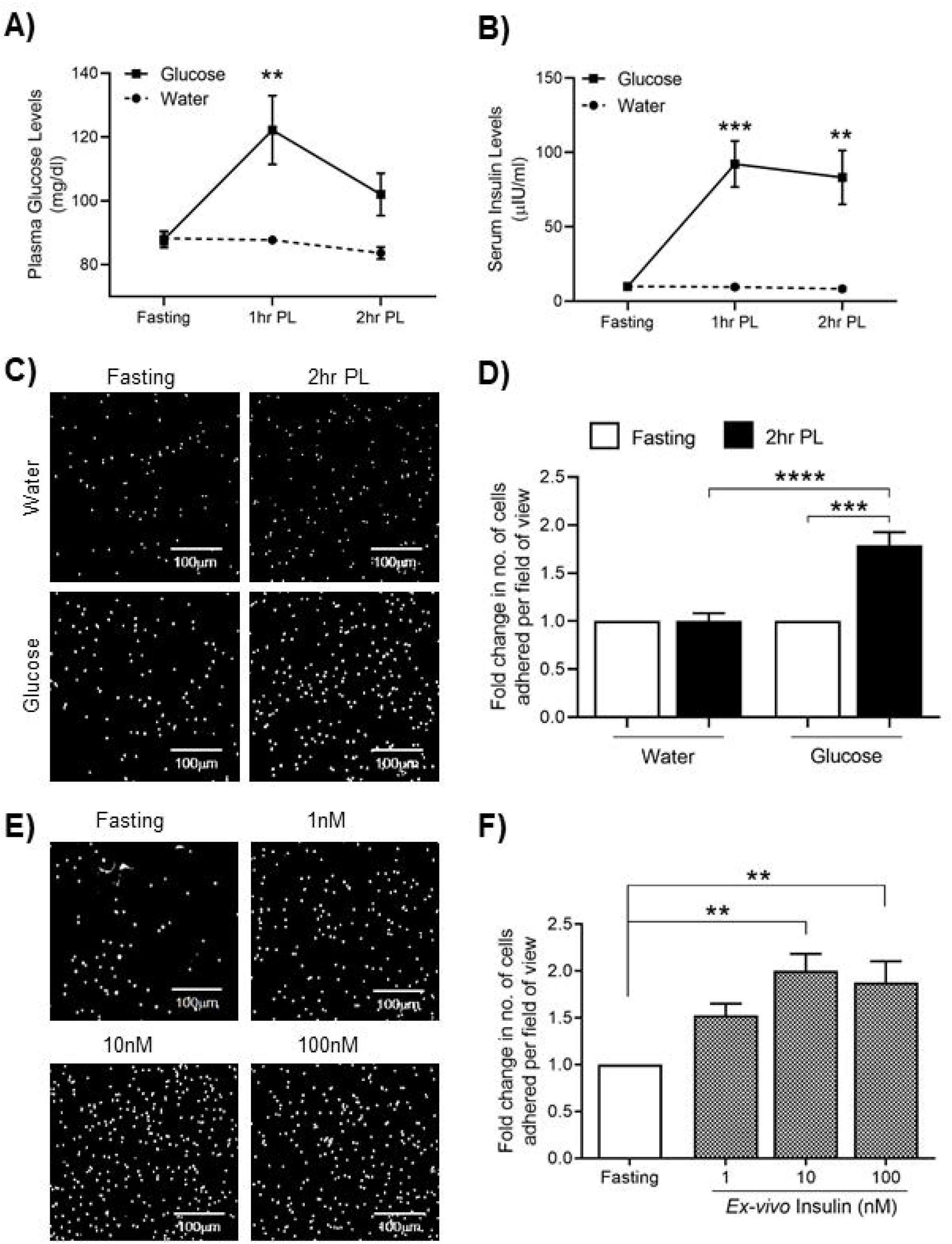
Enhanced adhesion of human PBMCs to fibronectin upon oral glucose load or *ex vivo* insulin treatment. A&B) Graphs showing the plasma glucose (n=10) and serum insulin levels (n=9) at fasting, 1-hour and 2-hour post water or glucose load in healthy individuals. ** p<0.01 and *** p<0.001 compared to fasting in paired two-tailed Student’s t-test. C) Representative images and D) bar graph, summarizing the number of PBMCs adhered to fibronectin (25µg/mL) for fasting and 2-hour post water or glucose load PBMCs of healthy subjects (n=10). *** p<0.001 and **** p<0.0001 in paired and unpaired two-tailed Student’s t-test, respectively. Scale bars correspond to 100 μm. PL denotes post load. E) Representative images and F) bar graph, summarizing the effect of *ex-vivo* insulin treatment of fasting PBMCs isolated from healthy NGT volunteers (n=4). ** p<0.01 compared to fasting in one-way ANOVA, Dunnett’s multiple comparison.

### 2.3. Oral glucose load enhances the adhesion of human PBMCs to fibronectin

Transmigration of circulating leukocytes across the basement membrane of blood vessels during diapedesis is necessary for their trafficking to the site of tissue damage or infection. Freshly isolated PBMCs from recruited subjects at fasting and at 2-hour post water or 2-hour post glucose load showed 100% staining for CD45, indicating a pure leukocyte population with no platelet contamination (Supplementary fig. 1B). These PBMCs were assayed for their ability to adhere to extracellular matrix proteins (ECM), fibronectin, collagen and gelatin. For the adhesion assay, equal number of PBMCs from all these conditions were seeded onto wells coated with ECM proteins and were incubated for 3 hours as described in the methods section. At the end of incubation, non-adherent cells were washed and adhered cells were fixed with paraformaldehyde and stained with DAPI for visualization. Oral glucose load specifically enhanced the adhesion of 2-hour post glucose load PBMCs to fibronectin by two-fold compared to fasting PBMCs (Fig. 1C and 1D). However, for the same subjects, there was no difference in fibronectin adhesion for 2-hour post water samples, indicating that this is a glucose-load specific phenomenon. Adhesion of PBMCs to other ECM proteins, collagen or gelatin (Supplementary fig. 1C), was not altered either upon water or upon glucose load in study subjects, demonstrating that an increase in adhesion in response to oral glucose load is specific to fibronectin. These observed differences in fibronectin adherence were not due to apoptosis or differences in viability of PBMCs across conditions either for water or for glucose load in study subjects as indicated by comparable 7-AAD and Annexin-V staining across different samples (Supplementary fig. 1D and 1E).

To assess whether increased fibronectin adherence influences the basal migration of PBMCs isolated from NGT subjects, cells were seeded onto fibronectin coated trans-well inserts, followed by measurement of migrated cells after 16 hours of incubation. Interestingly, despite the difference in fibronectin adhesion seen at 3 hours of adherence, both 2-hour post water and 2-hour post glucose load PBMC samples showed a comparable increase in basal migration in the trans-well assay (Supplementary fig. 2A and 2B). This may be explained by the difference in the incubation times in the adhesion versus the trans-well migration assays (i.e., 3 versus 16 hours). Basal migration here refers to migration of seeded PBMCs through the trans-well filters in absence of any agonists. Hence, results presented here indicate that physiological responses following glucose ingestion elicit adhesive changes in circulating leukocytes, possibly to modulate their immune, angiogenic and/or extra-vasation properties.

### 2.4. *Ex-vivo* treatment with insulin enhances the adhesion of fasting PBMCs to fibronectin

The most salient physiological response to increase in blood glucose levels is the consequent increase in circulating insulin levels. Biochemical data showed that at 2-hour post glucose load time point, while the glucose levels were back to normal range, insulin levels were still significantly higher compared to fasting in NGT subjects (Fig. 1A and 1B). This prompted us to determine if indeed *ex vivo* insulin treatment can enhance the fibronectin adherence of PBMCs similar to that observed for 2-hour post glucose load PBMC samples. In order to test the same, we isolated PBMCs from fasting samples of four healthy NGT individuals, and treated them *ex-vivo* with varying concentrations of endotoxin-free insulin for 2 hours. Treatment with insulin significantly increased the adhesion of fasting PBMCs to fibronectin (Fig. 1E and 1F). These results indicate that oral glucose load enhances the adhesion of human PBMCs to fibronectin possibly due to increased circulating levels of insulin. We employed 10nM insulin for the ensuing experiments in the study. It is noteworthy that none of the subjects whose samples were taken for the above experiments had any condition like infection, cancer or diabetes that would affect the activation status of immune cells. Also, all the experiments were performed with freshly isolated PBMCs, with no treatment whatsoever, other than insulin. Hence, the observed increase in fibronectin adhesion in response to insulin is for naïve PBMCs.

### 2.5. Oral glucose load as well as *ex-vivo* insulin treatment enhances adhesion of mice PBMCs to fibronectin

In order to better understand the molecular details of the observed phenomenon we set out to determine if insulin elicits similar effects with regard to fibronectin adherence in mice. We performed adhesion assays with PBMCs isolated from control and single high-dose streptozotocin (STZ) injected mice subjected to oral glucose load. Streptozotocin is particularly toxic to insulin producing pancreatic beta cells [30]. Single high-dose injection of STZ was chosen since it decreases the circulating insulin to non-detectable levels within 72 hours in multiple mice strains including C57BL/6 [31, 32]. Thus, it generates an insulin deficient mice model. Healthy control mice were subjected to intra-peritoneal injection of corresponding volume of citrate buffer. For the STZ group, only those mice with random blood glucose levels greater than 300 mg/dl post 72 hours of STZ injection confirming insulin loss were included in the study. Healthy control animals had mean random glucose levels less than 200 mg/dl (Supplementary fig. 2C). When OGTT was performed in these animals, both groups showed similar fasting glucose levels around 150mg/dl. However, 1-hour after oral glucose load, while control animals had glucose levels around 300mg/dl, STZ injected animals had even higher glucose levels at 1 hour compared to control animals (Supplementary fig. 2D).

PBMC samples isolated from mice for adhesion studies also showed that 98 percent cells stained positive for CD45, indicating a pure leukocyte population with no platelet contamination (Supplementary fig. 2E). PBMCs at fasting and 1-hour post load time points from normal and STZ injected mice were seeded onto fibronectin coated wells and incubated for 3 hours. After washing off non-adherent cells, the adhered cells were fixed and stained with DAPI for imaging. As can be seen from figure 2A and 2B, PBMCs isolated at 1 hour-post oral glucose load showed enhanced adhesion to fibronectin compared to fasting for control mice. In contrast, the adhesion of 1 hour post oral glucose load PBMCs was attenuated compared to fasting PBMCs from respective STZ treated animals, thereby alluding to the involvement of insulin.

**Figure 2:**
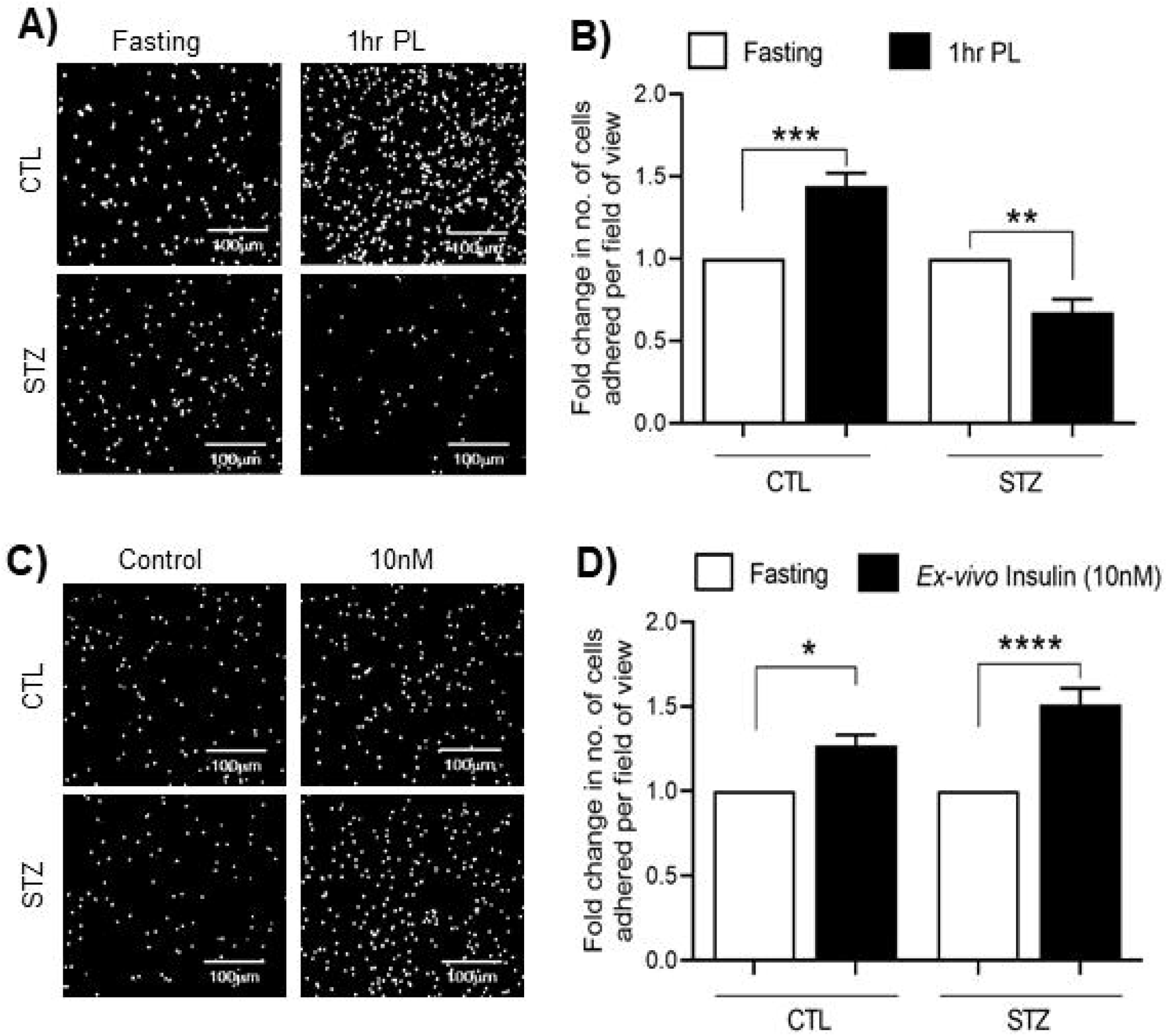
Enhanced adhesion of mice PBMCs to fibronectin upon oral glucose load or *ex vivo* insulin treatment. A) Representative images and B) bar graph, summarizing the fibronectin adhesion of fasting and 1-hour post oral glucose load PBMCs from control (n=10) and streptozotocin injected (n=12) mice. **p<0.01, *** p<0.001 in paired two-tailed Student’s t-test. C) Representative images and D) bar graph, summarizing the effect of *ex-vivo* insulin treatment of fasting PBMCs isolated from control (n=7) and streptozotocin injected mice (n=8). * p<0.05 and **** p<0.0001compared to respective fasting in one-way ANOVA with Tukey’s multiple comparison. Scale bars correspond to 100 μm.

We then set out to determine, if indeed insulin is responsible for observed adhesive effects even in mice similar to that observed for human PBMCs. For this fasting PBMCs from control and STZ mice were treated *ex-vivo* with 10nM insulin for 1-hour and were then seeded onto fibronectin coated wells. As seen in figure 2C and 2D, insulin treatment enhances fibronectin adherence of fasting PBMCs isolated from control as well as STZ treated mice. Similar to what was observed for human cells, fasting PBMCs, isolated from control mice showed enhanced adhesion to fibronectin upon *ex-vivo* treatment with insulin. Interestingly, fasting PBMCs isolated from the STZ-mice also showed increased adhesion to fibronectin following *ex-vivo* insulin treatment, confirming the role of insulin in enhancing the adherence of PBMCs to fibronectin. Thus, physiological increase in insulin following glucose load seems to be necessary for enhanced adherence of PBMCs to fibronectin even in mice just like that seen for humans.

### 2.6 Post glucose load PBMCs effectively home to injured vessels

Interaction of circulating leukocytes to ECM proteins such as fibronectin, plays a crucial role in homing and trafficking of immune cells to injured blood vessels and secondary lymphoid organs for tissue repair and immunity [33–36]. In order to determine if the post load PBMCs home to injured blood vessel better than fasting PBMCs, we performed intra-vital microscopy. For this, healthy C57BL/6J donor mice were subjected to oral glucose load. PBMCs isolated from these donor mice at fasting and 1 hour post glucose load were labelled with cell tracker dye Calcein-red orange as described in the methods section (Fig. 3A). Equal number of labelled fasting and post glucose load donor PBMCs were injected into corresponding healthy recipient mice through retro-orbital plexus. Homing of these labelled donor PBMCs to the endothelium devoid, exposed basement membrane of the mesenteric arteries of the recipient mice was then visualized through intra-vital microscopy. As seen in figure 3B and 3C, indeed the percent area of exposed basement membrane covered with adhered PBMCs was higher for 1 hour post glucose load donor PBMCs as compared to those with fasting donor PBMCs, demonstrating an enhanced homing of post-load PBMCs compared to fasting PBMCs. Thus, the oral glucose load mediated adhesive effects are seen even under flow conditions in an *in vivo* set-up wherein it assists in better homing of mononuclear cells to injured blood vessels with exposed basement membrane.

**Figure 3:**
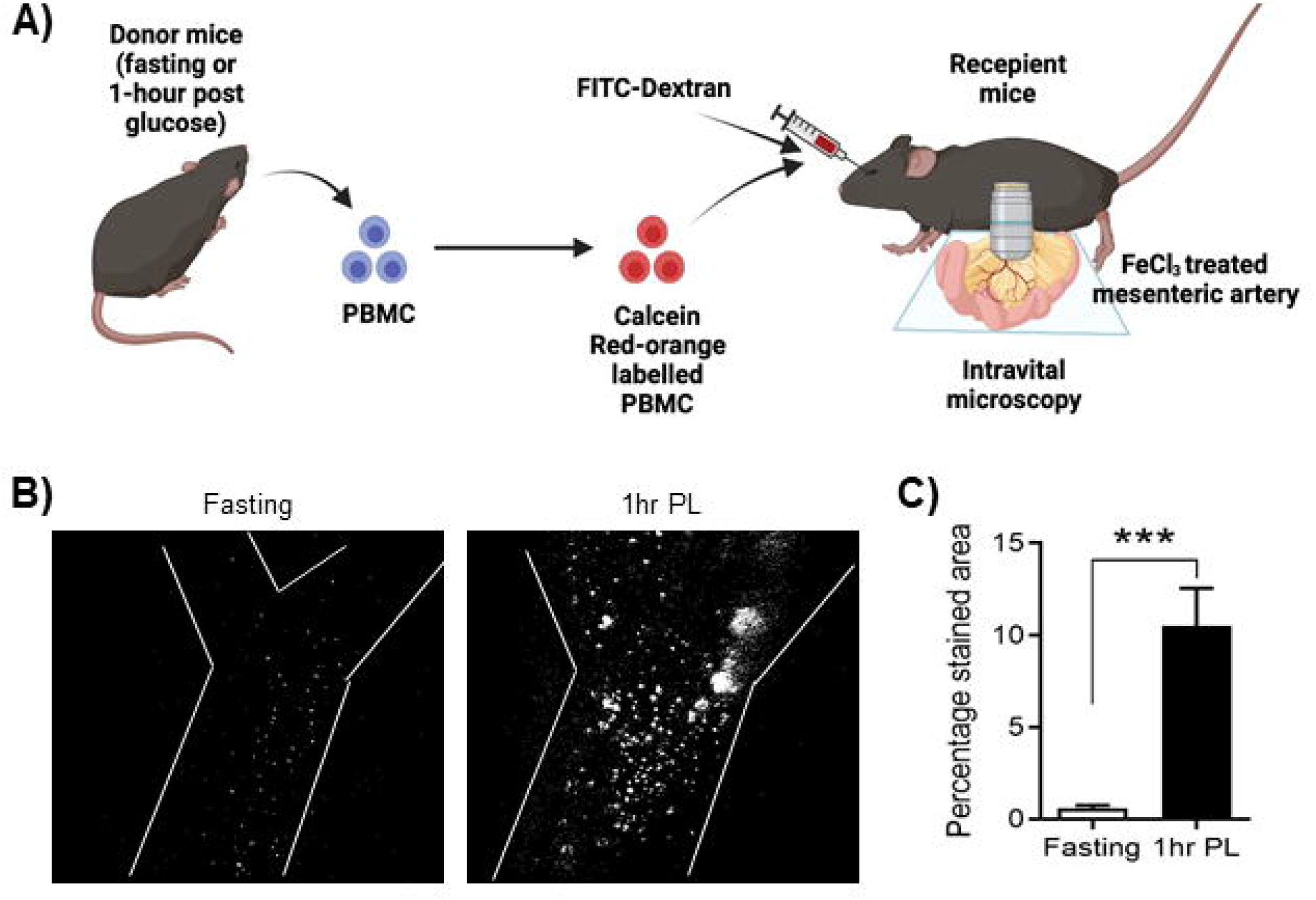
Enhanced adhesion of post-glucose load donor PBMCs to mesenteric artery of recipient mice. A) Cartoon depicting the methodology of intravital microscopy of PBMC adhesion to the denuded mesenteric artery in live recipient mice. B) Representative images and C) bar graph, summarizing the intra-vital imaging of the adhesion of fasting and 1-hour post glucose load donor mice PBMCs to denuded mesenteric arteries of recipient mice for eight independent animals. ***p<0.001 in unpaired two-tailed Student’s t-test.

### 2.7. Insulin enhances lymphocyte adhesion via IGF-1R auto-phosphorylation

Human PBMCs majorly contain lymphocytes (around 80-90%) and monocytes. Within lymphocytes, T-cells are the major constituents (around 80%) [37]. Flow-cytometry analysis of isolated human PBMC samples confirmed that they were largely comprised of lymphocytes as shown by CD5 surface expression (Supplementary fig. 3A to 3E). Hence, we sought to see if insulin treatment can augment the adhesion of lymphocytes to fibronectin. A time-course treatment of serum starved Jurkat T-lymphocytes with 10nM insulin indeed induced a significant increase in fibronectin adhesion as early as 1 minute, which was sustained till 120 minutes, compared to respective time controls of untreated cells (Fig. 4A and 4B). It is important to note that Jurkat cells were not activated with T-cell receptor agonists prior to insulin treatment or during incubation on fibronectin coated wells in any of these experiments. Hence, the phenomena observed are for naïve, resting lymphocytes. Since Jurkat T-cells recapitulated the phenomenon found in primary PBMCs, they were further used in cell culture experiments to delineate the molecular mechanism of insulin-induced increase in fibronectin adhesion.

**Figure 4:**
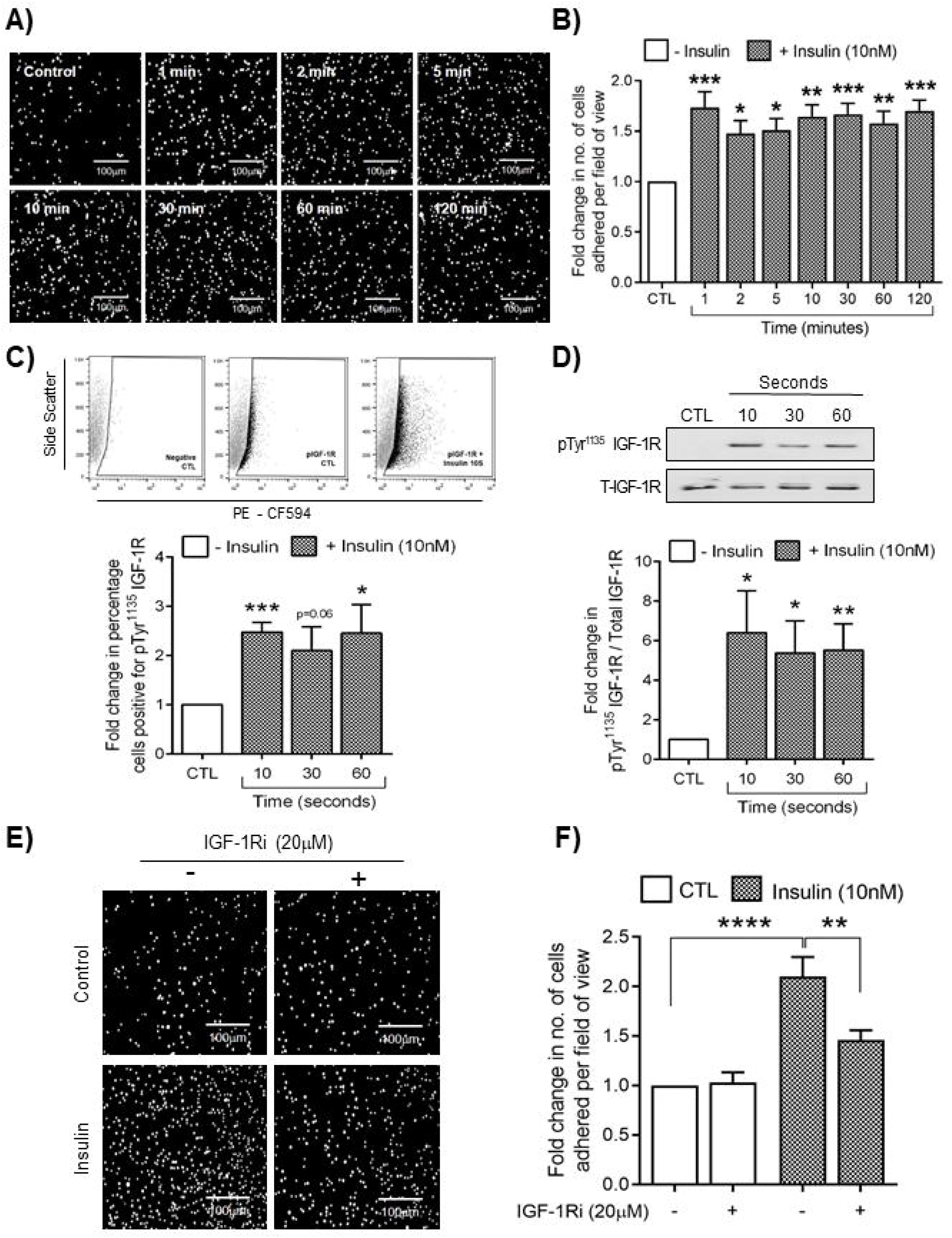
Insulin enhances fibronectin adherence of Jurkat T-cells through IGF-1 receptor phosphorylation. A) Representative images and B) bar graph, summarizing the effect of time-course treatment of serum starved Jurkat cells with 10nM insulin from nine independent experiments. *p<0.05, **p<0.01 and ***p<0.001, in one-way ANOVA with Dunnett’s multiple comparison. C) Dot plots depicting the cells positive for phospho-Tyr^1135^ IGF-1R in control and insulin treated Jurkat cells and bar graph, summarizing the fold change in the percentage of gated cells positive for pIGF-1R following insulin treatment in serum starved Jurkat cells from four independent experiments. *p<0.05 and ***p<0.001, in unpaired two-tailed Student’s t-test. D) Representative Western blot images and bar graph summarizing the levels of phosphorylation at Tyr^1135^ of IGF-1 receptor during a time course treatment of Jurkat cells with 10nM insulin, from six independent experiments. *p<0.05 and **P<0.01 in unpaired two-tailed Student’s t-test. E) Representative images and F) bar graph, summarizing the effect of inhibition of IGF-1 receptor auto-phosphorylation on insulin-induced adhesion of Jurkat cells to fibronectin for seven independent experiments. **p<0.01 and ****p<0.0001 in one-way ANOVA with Tukey’s multiple comparison. Scale bars correspond to 100 μm.

It is reported that insulin receptors are not expressed on the surface of resting lymphocytes [27]. In order to confirm the same with our cell culture model, we performed surface flow-cytometry to detect the presence of insulin receptors (IR) on Jurkat T-cells. As can be seen from Supplementary fig. 3F, insulin receptors were not expressed on the surface of serum starved Jurkat T-cells, as opposed to HEK293 cells, which expressed IRs on their surface. Given that insulin can also bind and activate IGF-1 receptor (IGF-1R), which are known to be expressed on naïve T-cells [38], we performed flow-cytometry and Western blotting experiments to detect the phosphorylation of IGF-1R in serum starved Jurkat cells at the Tyr^1135^ residue, following insulin treatment. There was a significant increase in both the percentage of cells positive for phospho-IGF-1R (pIGF-1R) (flow-cytometry) and the overall amount of phosphorylation of the IGF-1 receptor (Western blotting) upon insulin treatment, as early as 10 seconds which was sustained till 1 minute (Fig. 4C and 4D). Further, inhibiting the auto-phosphorylation of IGF-1R through pre-treatment of serum starved Jurkat cells with 20µM PQ401, significantly attenuated the insulin-induced adhesion of lymphocytes to fibronectin (Fig. 4E and 4F). These results show that insulin-induced increase in fibronectin adhesion is mediated in part through IGF-1R auto-phosphorylation in response to insulin (10nM), as early as 10 seconds in resting Jurkat lymphocytes.

### 2.8. Insulin activates PLC-γ1 to mediate fibronectin adherence of lymphocytes

We looked at the phosphorylation status of various intra-cellular intermediates of insulin signalling in lymphocytes in a time-course experiment with 10nM insulin. Insulin is classically known to mediate the activation of two major pathways, namely: the PI-3Kinase/Akt axis and the mitogen activated protein kinase pathway (MAPK). While p38-mitogen activated protein kinase (p38MAPK), and the extracellular-signal regulate kinase (ERK) did not show any change (data not shown) upon insulin treatment, Ser^473^ phosphorylation of Akt was observed from 5 minutes until 30 minutes of insulin treatment before returning to control levels by 1 hour (Fig. 5A and 5B). Since, changes in intra-and extra-cellular concentrations of calcium influence cellular adhesion we set out to determine if insulin can activate phospholipase C (PLC). Intriguingly, insulin enhanced the phosphorylation of PLC-gamma 1 (PLC-γ1) at Tyr^783^ as early as 10 seconds, with significant increase at 30-second, 5-minute, and 10-minute time points. Tyr^783^ phosphorylation levels returned to normal by 60 minutes, when compared with respective time control (Fig. 5A and 5B). Given that insulin elicits activation of IGF-1R as early as 10 seconds, we next determined the effect of IGF-1R inhibition on insulin induced PLC-γ1 phosphorylation in Jurkat cells. Inhibition of IGF-1R auto-phosphorylation with 20μM PQ401, blocked PLC-γ1 phosphorylation (Fig. 5C). This indicates that PLC-γ1 is down-stream of IGF-1R signalling. Moreover, inhibition of PLC activation by pre-treatment of serum starved Jurkat cells with 1µM U73122 (PLC inhibitor) for 1 hour abrogated insulin-induced (10nM, 10 minutes) increase in fibronectin adhesion (Fig. 5E and 5F), confirming the involvement of PLC-γ1 in the phenomenon. These results show that PLC-γ1, which is down-stream of IGF-1R mediates the insulin-induced fibronectin adherence of lymphocytes.

**Figure 5:**
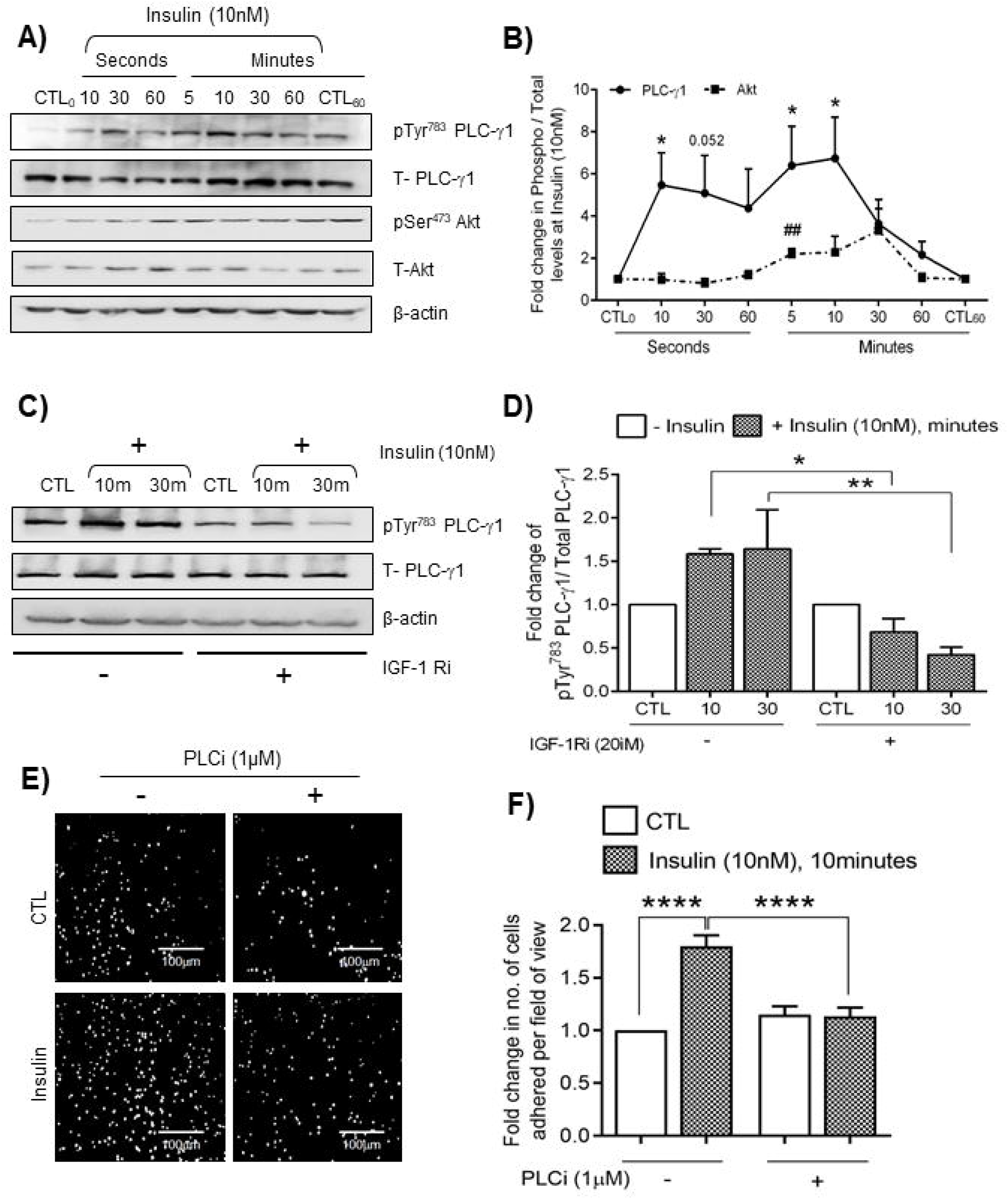
**Insulin-induced increase in fibronectin adhesion of Jurkat T-cells is dependent on Tyr^783^ phosphorylation of PLC-**γ**1.** A) Representative Western blot images and B) graph, summarizing the levels of phosphorylation at Tyr^783^ of PLC**-**γ1 (n=5 independent experiments) and at Ser^473^ of Akt (n=3 independent experiments), during a time course treatment of Jurkat cells with 10nM insulin for respective time points. *p<0.05 and ##<0.01, of a given time point, compared to respective control in unpaired two-tailed Student’s t-test. C) Representative Western blot images and D) bar graph, summarizing the levels of phosphorylation at Tyr^783^ of PLC**-**γ1 (n=5 independent experiments), upon treatment with 10nM insulin, with or without the inhibition of IGF-1 receptor auto-phosphorylation. *p<0.05 and **p<0.01 in one-way ANOVA with Tukey’s multiple comparison. E) Representative images and F) bar graph, summarizing the effect of PLC inhibition on insulin-induced adhesion of Jurkat cells to fibronectin from six independent experiments. ****p<0.0001 in one-way ANOVA with Tukey’s multiple comparison. Scale bars correspond to 100 μm.

Considering that PLC activation leads to intracellular calcium rise which along with extracellular cations, Mn^2+^ and Mg^2+^, plays a crucial role in integrin mediated cellular adhesion [39, 40], we next determined the role of calcium in insulin mediated fibronectin adherence. Pre-treating Jurkat cells with intracellular calcium chelator BAPTA-AM, inhibited the insulin-induced increase in lymphocyte adhesion to fibronectin (Supplementary fig. 4A and 4B). Chelating the extracellular divalent cations with EDTA also showed similar inhibition (Supplementary fig. 4C and 4D) thereby, highlighting the involvement of intracellular as well as extracellular divalent cations in insulin mediated fibronectin adherence of lymphocytes.

### 2.9. Enhanced lymphocyte adhesion to fibronectin is due to activation of integrin β1

Interaction of cells with extra-cellular matrix proteins is mediated by integrins expressed on the cell surface. We performed flow-cytometry on fasting human PBMCs (n=5) and serum starved Jurkat T-cells (n=5) to assess the expression of various integrins on their surface. Integrins α4, α5, αL, β1 and β2 were majorly expressed on the surface of both cell types i.e., PBMCs and Jurkat T cells (Supplementary fig. 5A). αX was expressed in a small proportion of both PBMCs and Jurkat. A minuscule proportion of PBMCs also expressed integrins αE, αM, and αX, these were however absent in Jurkat T-cells. This was expected, considering PBMCs are a heterogeneous cell mixture while Jurkat T-cells are pure lymphocytes. Integrin αVβ3 was expressed more on Jurkat T-cells than in PBMCs (Supplementary fig. 5A).

Integrin mediated adhesion of cells can be enhanced by an increase in their surface expression or by change in their conformation leading to activation [41]. We compared the surface expression of α4, α5, αL, β1 and β2 integrins on fasting and 2-hour post glucose load PBMCs obtained from NGT subjects and did not observe any change either in the percentage of cells positive or in the surface expression of integrins as assessed through mean fluorescence intensity (Supplementary fig. 5B and 5C). Integrins α4 or α5 associate with β1 and αL associates with β2 to form functional integrin complexes VLA4, VLA5 and LFA1 respectively. Hence, we performed flow-cytometry, using antibody clones TS2/16 and m24 that specifically identify the activated conformations of β1 and β2 integrins respectively. While there was no change in the activation status of integrin β2 (Supplementary fig. 5D), activation of integrin β1 increased following insulin treatment in serum starved Jurkat cells (Fig. 6A and 6B). Treatment of cells with 5μM BIO5192, an inhibitor which binds to α4β1 integrin and prevents its interaction with ligands, attenuated the insulin-induced adhesion (10nM, 10 minutes) of Jurkat lymphocytes to fibronectin, thus confirming its involvement in the phenomenon (Fig. 6C and 6D). Further, inhibiting IGF-1R auto-phosphorylation by pretreatment of serum starved Jurkat cells with 20μM PQ401 prevented the insulin-induced (10nM) activation of integrin β1 as shown by activation specific antibody (TS2/16) binding in flow-cytometry (Fig. 6E and 6F). These results show that insulin-induced increase in fibronectin adherence of lymphocytes is a result of IGF-1 receptor mediated PLC-γ1 activation leading to inside-out activation of integrin β1 on the cell surface as summarized in the graphical abstract.

**Figure 6:**
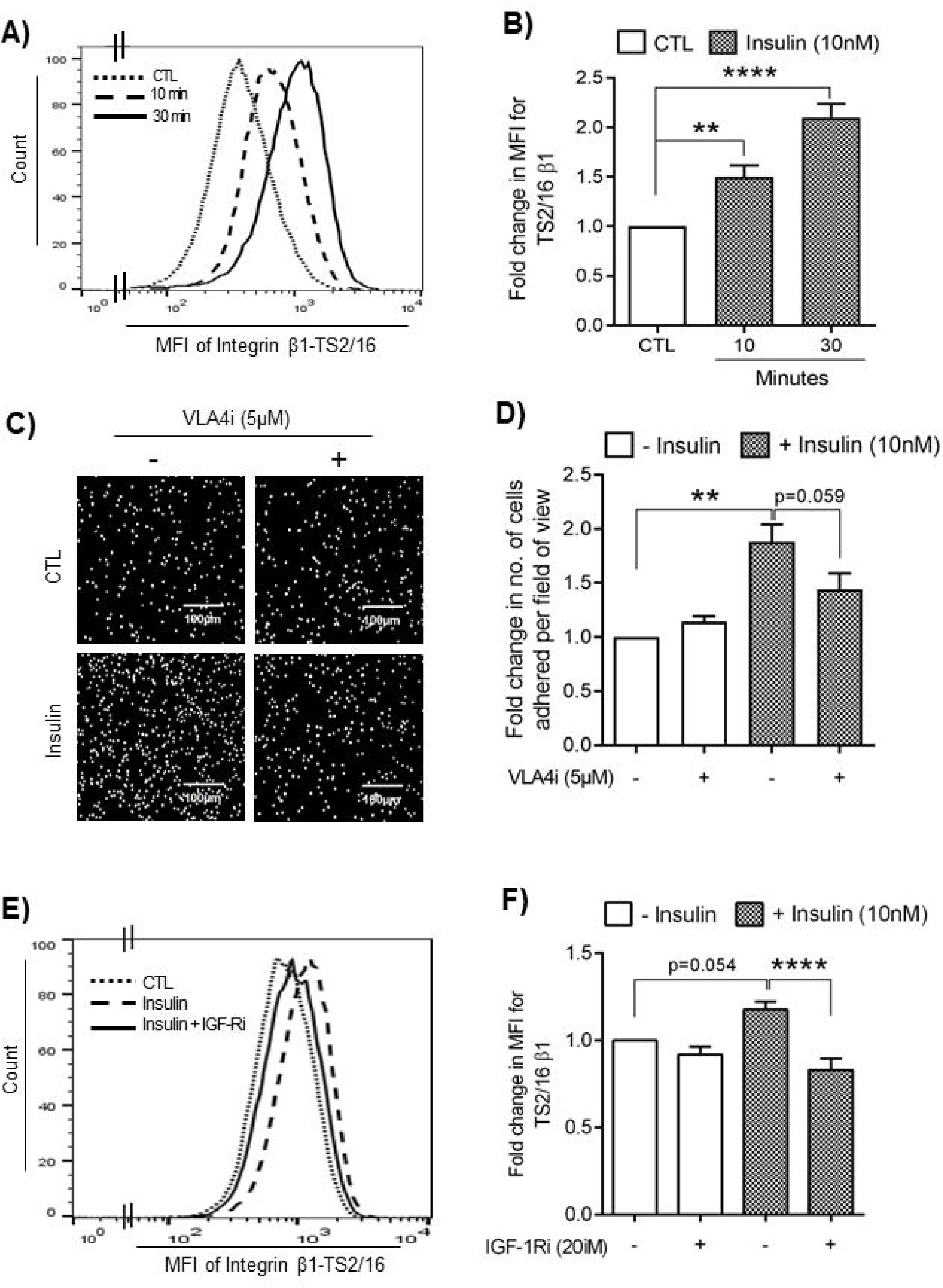
Insulin enhances adhesion of Jurkat T-cells to fibronectin through activation of integrin. β1. A) Representative histograms and B) bar graph, summarizing the activation status of integrin β1 in Jurkat cells upon treatment with 10nM insulin, as shown by mean fluorescence intensity (MFI) due to binding of activation specific antibody clone TS2/16, from five independent experiments. **p<0.01 and ****p<0.0001, compared to control in one-way ANOVA with Dunnett’s multiple comparison. C) Representative images and D) bar graph, summarizing the effect of integrin β1 inhibition on insulin-induced fibronectin adhesion of Jurkat cells from five independent experiments. **p<0.01 in one-way ANOVA with Tukey’s multiple comparison. E) Representative histograms and F) bar graph, summarizing the effect of inhibition of IGF-1 receptor auto-phosphorylation on insulin-induced integrin β1 activation, from seven independent experiments. ***p<0.001 in one-way ANOVA with Tukey’s multiple comparison.

### 3. Discussion

The role of insulin signaling in different subsets of leukocytes which may or may not express the insulin receptors (IR), is now beginning to emerge. It was demonstrated that indeed insulin elicits intracellular signaling in human leukocytes, however it is unclear as to what is the functional relevance of this signaling, despite it being disrupted in diabetic patients [42, 43]. Others have shown that insulin primes activated neutrophils toward phagocytosis [1]. However, the physiological role of insulin signaling, if any, in quiescent circulating lymphocytes remains elusive. Of note, the expression of insulin receptors on the surface of T-lymphocytes is detected only upon antigen presentation or mitogen activation [2, 27]. In the activated T-cells, insulin signaling promotes nutrient uptake, glycolysis, proliferation, cytokine production, differentiation and cytotoxic responses [2]. In the present study, we demonstrate that naïve PBMCs from humans and mice, as well as T-cells, respond to insulin to promote fibronectin adherence, and this involves IGF-1R mediated PLCγ-1 phosphorylation leading to activation of β1 integrin (Graphical abstract). To the best of our knowledge, this is the first study to identify a potentially relevant role of post-prandial insulin spikes on the physiology of circulating but naïve T-cells during feeding as seen in this study in response to oral glucose load.

Impact of daily fasting and feeding cycles on circulating immune cells is currently unknown. Consumption of carbohydrate rich diet or oral glucose load elicit post-prandial increase in insulin, incretins and gut-microbiota derived metabolites [44, 45]. Since circulating leukocytes are constantly exposed to oscillations in blood biochemistry, they are bound to be the early responders to these endocrine changes. It is worthwhile to note that almost all fractions of PBMC including resting T-cells and Jurkat T cells do express IGF-1 receptors, and intriguingly, the surface expression of IGF-1 receptors decreases upon T cell activation [46–48]. As insulin binds to the IGF-1R with lower affinity [46], one may argue that the observed effects are due to circulating IGF-1 levels. It should however be noted that multiple studies have demonstrated that unlike insulin, the circulating levels of bioactive IGF-1, decline following oral glucose load [49–51]. Alternatively, one may argue that the increase in fibronectin adherence post glucose load, is due to the direct effect of glucose dependent insulinotropic peptide (GIP) and glucagon like peptide-1 (GLP-1) on lymphocytes [44]. Although we cannot negate this possibility, but, given that incretins themselves enhance insulin secretion and since the adhesive effects of post-load PBMCs were abrogated in insulin deficient streptozotocin mice, we propose that insulin by virtue of inducing IGF-1R auto-phosphorylation is predominantly responsible for the observed phenomenon in this study.

High affinity cryptic insulin receptors (IR) are detected on the surface of T-cells only upon their activation [27], unlike metabolic tissues and monocytes which express them constitutively. Instead, naïve T-cells exhibit surface expression of IGF-1R, which decreases upon their activation [48]. In the current study, even in the absence of an activating stimulus, insulin promoted IGF-1R auto-phosphorylation and intracellular signalling in naïve lymphocytes. Canonical insulin signalling involves auto-phosphorylation of insulin receptor followed by phosphorylation of IRS-1 and activation of PI-3 Kinase and Akt axis. Although insulin treatment did lead to Ser^473^ phosphorylation of Akt in Jurkat-cells at 5 minutes, the adhesive effects of insulin were seen as early as one minute. Also, the earliest signalling events triggered by insulin were IGF-1R auto-phosphorylation and Tyr^783^ phosphorylation of PLC-γ1, which were seen as early as 10 seconds. PLC-γ1 has been shown to physically interact with phosphorylated receptor tyrosine kinases through its SH-2 and PH domains [52]. This could explain the phosphorylation of PLC-γ1 immediately following the auto-phosphorylation blocked insulin mediated PLC-γ1 activation, β1 integrin activation and consequently fibronectin adherence. Thus, insulin led IGF-1R auto-phosphorylation as early as 10 seconds, along with concurrent activation of PLC-γ1 yielding to enhanced fibronectin adherence, indicates an inside-out activation of integrin β1 in circulating PBMCs and naïve lymphocytes.

Functions such as adhesion, migration, endothelial interaction and extravasation for naïve T-cells are majorly regulated by integrin β1, and these are subsequently taken over by integrin β2 upon activation in T-cells [53]. PLC-γ1 is a key player in promoting both ‘Outside-in’ and ‘Inside-out’ modes of integrin signalling. In fact, PLC-γ1 can directly interact with the cytoplasmic tail of integrin β1 to induce a conformational change for integrin activation [54]. Both integrin β1 and PLC-γ1 are highly expressed in circulating T-lymphocytes and chimeric mice generated from their respective null stem cells allude to their involvement in hematopoiesis [55, 56]. Thus, apart from playing a critical role in TCR mediated activation of integrin β2 (LFA-1) [57], the current study identifies a novel non-effector function of PLC-γ1 in insulin mediated fibronectin adherence of naïve T-cells.

Fibronectin binding integrins VLA-4 (α4β1), VLA-5 (α5β1) and LFA-1 (αLβ2) in leukocytes are indispensable for immune cell trafficking, activation and paracrine activity [33–36]. Rolling, adhesion and trans-migration of leukocytes to and through the vascular endothelium involves temporal activation of different subsets of integrins [58]. Even periodic circulation and recirculation of naïve T-cells through blood into the lymphoid and non-lymphoid organs, and back, via lymph, involves integrin dependent interaction of leukocytes with the extracellular matrix (ECM) proteins, as seen in the high endothelial venules (HEVs) of lymph nodes [36, 59]. Similarly, β1 integrin expressing lymphocytes home into multiple non-lymphoid organs be it the skin, lungs or the brain, wherein, their interaction with ECM assists in T-cell receptor signaling [36,60,61]. Furthermore, presence of β1 integrin at the immune synapse between a T-cell and an antigen presenting cell (APC) particularly in non-lymphoid organs assists in local immunity [62, 63]. Hence, leukocyte integrins are necessary for their trafficking and immune response. It is worth noting that in the quiescent state, the integrins present on naïve T-cells have lower affinity and avidity specifically towards fibronectin [34], however, binding of homing cytokines such as SDF-1α to their cognate receptors, elicits an ‘inside-out’ activation of these integrins (α4β1) to enhance their fibronectin binding [64, 65]. In the current study we report that similar to homing cytokines, insulin also elicits fibronectin adherence of naïve circulating T-cells via activation of integrin β1. Based on these observations and our intra-vital microscopy data, it is fair to speculate that the responsiveness of circulating quiescent T-cells to post-prandial insulin spikes would contribute to their periodic homing and recruitment due to enhanced fibronectin adherence, to lymphoid as well as non-lymphoid organs for routine surveillance, repair and localized immunity. Oral glucose load in healthy individuals also regulates the effector gene expression in innate and adaptive immune cells [8, 26]. Studies have demonstrated that β1 integrins are necessary for the paracrine activity of T-cells including Jurkat T-cells, for expression and release of IFN-γ and VEGF [66]. Hence, it is also likely that increased fibronectin adherence of circulating T-cells in the absence of APCs may contribute to local angiogenesis and immunity in tissues through release of these factors upon homing. Thus, our study points to a tissue homeostatic role of circulating, insulin responsive naïve T-cells even in absence of antigen presentation.

It is imperative that the suggested adhesive effects of post-prandial insulin spikes on circulating cells are stringently regulated. Any alterations, either in cellular signaling or in the kinetics, magnitude, duration and/or the nature of the post-prandial responses will have detrimental consequences. For instance, persistent expression, activation and interaction of VLA-4 with fibronectin, perpetuates build-up of immune cells in the joints of rheumatoid arthritis patients [67]. Similarly, excessive infiltration of leukocytes into the adipose and hepatic tissues is a hall-mark feature of insulin resistance and diabetes [68, 69]. Likewise, enhanced expression of β1 integrin is noted in multiple cancers, where it is associated with chemo-resistance, radio-resistance, metastasis, tumor angiogenesis and reduced survival [70–77]. Interestingly, it is the activation of β1 integrin that triggers the transition from dormancy to metastasis by aiding in the cross talk of dissociated metastatic cancer cells with the tumor micro-environment [75, 76]. Unfortunately, VLA-4 mediated tumor invasion of immunosuppressive myeloid cells and Tregs also contributes to immune evasion in cancer [78]. Given the fact that over-feeding induced blood changes such as hyperinsulinemia and diabetes, are associated with cancer progression and blunting of chemotherapy [79, 80], our data of oral glucose load induced activation of β1 integrin and consequent fibronectin adherence, sheds light on the likely molecular mechanism of chemo-resistance and metastasis observed during metabolic syndrome. It is thus important to study the cell specific effects of post-prandial insulin spikes on different subsets of immune cells in diabetic subjects to delineate the pathophysiology of associated comorbidities.

Our study has the following limitations: 1) Women were not included in this study. This is because, the hormonal changes during the menstrual cycle, even in healthy women influence circulating counts and subsets of immune cells [81]. Hence, a stand-alone study in women accounting for stage specific variations in immune cells during menstruation while performing OGTT is necessary. 2) Owing to the ethical constraints imposed on the amount of blood sample that could be collected, we could not examine the effects of OGTT or insulin treatment on specific lymphocyte subsets and changes in their functional properties. 3) Since this study was carried out only in Asian Indian men, similar studies in other ethnicities is desirable. In conclusion, we demonstrate that oral glucose load in healthy men and mice enhances adherence of PBMCs to fibronectin. Further, as summarised in the graphical abstract, we demonstrate that insulin mediates lymphocyte adherence to fibronectin through a non-canonical signalling involving IGF-1R auto-phosphorylation, PLC-γ1 and integrin β1 activation, thereby identifying a non-metabolic role of insulin in quiescent naïve T-cells.

## Author contributions

MACT performed PBMC isolation, adhesion assays, flow-cytometry experiments, inhibitor experiments, analyzed and interpreted the data, performed statistical analyses and wrote the initial draft of the manuscript. SS performed Western blotting experiments, inhibitor experiments and analyzed the Western blotting data. MACT, HA and DP performed animal experiments. AAN performed adhesion imaging and statistical analysis. MMR coordinated human volunteer recruitment, screening, and sample collection. RB supervised and analysed animal experiments. KJ designed, supervised and interpreted intra-vital microscopy experiments. MKS provided intellectual input. MDX conceptualized the study, procured funding, interpreted the results, wrote and revised the manuscript for intellectual content.

## Acknowledgements

Authors acknowledge Sumeet Parihar (Pharmacology Division, CSIR-CDRI, Lucknow), the phlebotomists and the laboratory staff at the Institute Hospital, IIT Madras, Chennai. Authors thank also Dr. Vani Janakiraman (Department of Biotechnology, IIT Madras, Chennai) for critical reading of the manuscript and suggestions.

## Supplementary figure legends

**Supplementary figure 1:** A) Table summarizing the total and differential WBC counts in blood at fasting and 2-hour post water and oral glucose load in healthy NGT subjects (n=10). Total WBC count is shown in cells / mm^3^ and differential cell count in percentage. B) Bar graph, summarizing the purity of isolated human PBMC samples, at all conditions as indicated by surface staining for CD45 (n=8 subjects). C) Bar graph, summarizing the adhesion of PBMCs of fasting and 2-hour post water or glucose load samples of NGT subjects to collagen (n=10) and gelatin respectively (n=9). D) Bar graphs summarizing the percentage of live cells as indicated by negative staining for 7-AAD, and E) the percentage of early apoptotic cells as indicated by AnnexinV-PE staining within the 7AAD^-^ population in PBMCs across all conditions.

**Supplementary figure 2:** A) Representative images and B) bar graph, summarizing the basal migration of PBMCs, from fasting and 2-hour post water (n=10) or glucose load samples (n=8) from NGT subjects, through trans-well filters of 8 μm pore size. * indicates p<0.05 compared to respective fasting values in paired two-tailed Student’s t-test. C) Bar graph summarizing the random blood glucose levels in control (CTL, n=11) and streptozotocin injected mice (STZ, n=20), 72 hours after the injection. ****p<0.0001 in unpaired two-tailed Student’s t-test. D) Bar graph summarizing the blood glucose levels at fasting and 1-hour post oral glucose load in control (CTL, n=10) and streptozotocin injected mice (STZ, n=12). ***p<0.001 and ****p<0.0001 in paired and unpaired two-tailed Student’s t-test. E) Bar graph summarizing the purity of isolated mouse PBMC samples, at all conditions as indicated by surface staining for CD45.

**Supplementary figure 3:** A) Representative dot plot depicting the lymphocyte-monocyte (LM) gating. B) Representative dot plot demonstrating gating for CD5-FITC and CD14-PE negative cells within the LM gated cells. C&D) Percentage of cells positive for CD5-FITC and CD14-PE at fasting and at 2-hour post oral glucose load in human PBMC samples. E) Bar graph summarizing the proportion of lymphocytes and monocytes within the PBMCs of fasting and 2-hour post oral glucose load samples from study subjects (n=3). F) Histograms depicting surface expression of insulin receptor on serum starved Jurkat and HEK cells (n=3).

**Supplementary figure 4:** A) Representative images and B) bar graph summarizing the effect of chelation of intra-cellular calcium by BAPTA-AM (10µM) on insulin-induced fibronectin adhesion of Jurkat cells, from seven individual experiments. C) Representative images and D) bar graph, summarizing the effect of chelation of extra-cellular divalent cations by EDTA on insulin-induced increase in adhesion of Jurkat cells to fibronectin. *p<0.05, ***p<0.001 and ****p<0.0001 in one-way ANOVA with Tukey’s multiple comparison.

**Supplementary figure 5:** A) Table summarizing the percentage of cells positive for surface expression of respective integrins on fasting PBMCs from NGT subjects (n=5) and serum starved Jurkat cells (n=5). B) and C) Bar graphs summarizing the percentage of cells positive and mean fluorescence intensity respectively, of the surface expression of respective integrins on fasting and 2-hour post glucose load PBMCs from NGT subjects (n=5). D) Bar graph summarizing the activation status of integrin β2 in Jurkat cells upon treatment with 10nM insulin, as shown by mean fluorescence intensity due to binding of activation specific antibody clone m24 (n=6).

## STAR Methods

### KEY RESOURCES TABLE

STAR Methods: KEY RESOURCES TABLE:
| REAGENT or RESOURCE | SOURCE | IDENTIFIER |
| --- | --- | --- |
| **Antibodies** |  |  |
| Akt Antibody | Cell Signaling Technology | Cat# 9272<br>RRID: AB_329827 |
| Anti-Insulin Receptor (Ab-1) Mouse mAb (CII 25.3) | Merck-Millipore | Cat# GR05<br>RRID: AB_564749 |
| APC anti-human CD29 Antibody (Clone: TS2/16) | Biolegend | Cat# 303008<br>RRID: AB_314324 |
| APC anti-human CD11a/CD18 (LFA-1) Antibody (Clone: m24) | Biolegend | Cat# 363410<br>RRID: AB_2716070 |
| BD™ PerCP Mouse Anti-Human CD45 | BD Biosciences | Cat# 347464<br>RRID: AB_400307 |
| BD Pharmingen™ APC-Cy™7 Rat Anti-Mouse CD45 | BD Biosciences | Cat# 557659<br>RRID: AB_396774 |
| BD Pharmingen™ FITC Mouse Anti-Human CD5 | BD Biosciences | Cat# 555352<br>RRID: AB_395756 |
| BD™ PE Mouse Anti-Human CD14 | BD Biosciences | Cat# 347497<br>RRID: AB_400312 |
| BD Pharmingen™ APC- Mouse Anti-Human CD49d | BD Biosciences | Cat# 559881<br>RRID: AB_398681 |
| BD Pharmingen™ PE Mouse Anti-Human Integrin α5 chain (CD49e) | BD Biosciences | Cat# 555617<br>RRID: AB_395984 |
| BD Horizon™ BV786 Mouse Anti-Human CD29 | BD Biosciences | Cat# 564815<br>RRID: AB_2738970 |
| BD Horizon™ BV711 Mouse Anti-Human CD11a | BD Biosciences | Cat# 563935<br>RRID: AB_2738494 |
| BD Horizon™ BV421 Mouse Anti-Human CD18 | BD Biosciences | Cat# 562871<br>RRID: AB_2737855 |
| BD Horizon™ PE-CF594 Mouse Anti-Human CD11b | BD Biosciences | Cat# 562399<br>RRID: AB_2737613 |
| BD OptiBuild™ BV510 Mouse Anti-Human CD51/CD61 | BD Biosciences | Cat# 744089<br>RRID: AB_2741984 |
| BD Horizon™ BV650 Mouse Anti-Human CD11c | BD Biosciences | Cat# 563404<br>RRID: AB_2869490 |
| BD Horizon™ BB515 Mouse Anti-Human CD103 | BD Biosciences | Cat# 564578<br>RRID: AB_2738852 |
| BD™ PerCP Mouse IgG1, κ Isotype Control | BD Biosciences | Cat# 349044<br>RRID: AB_400399 |
| BD Pharmingen™ APC-Cy™7 Rat IgG2b κ Isotype Control | BD Biosciences | Cat# 552773<br>RRID: AB_394459 |
| BD Pharmingen™ APC Mouse IgG1 κ Isotype Control | BD Biosciences | Cat# 555751<br>RRID: AB_398613 |
| BD Pharmingen™ FITC Mouse IgG1, κ Isotype Control | BD Biosciences | Cat# 555748<br>RRID: AB_396090 |
| BD Pharmingen™ PE Mouse IgG2b κ Isotype Control | BD Biosciences | Cat# 555743<br>RRID: AB_396086 |
| BD Pharmingen™ PE Mouse IgG1, κ Isotype Control | BD Biosciences | Cat# 555749<br>RRID: AB_396091 |
| BD Horizon™ BV786 Mouse IgG1, κ Isotype Control | BD Biosciences | Cat# 563330<br>RRID: AB_2869484 |
| BD Horizon™ BV711 Mouse IgG1, κ Isotype Control | BD Biosciences | Cat# 563044<br>RRID: AB_2869449 |
| BD Horizon™ BV421 Mouse IgG1, κ Isotype Control | BD Biosciences | Cat# 562438<br>RRID: AB_11207319 |
| BD Horizon™ PE-CF594 Mouse IgG1, κ Isotype Control | BD Biosciences | Cat# 562292<br>RRID: AB_11207243 |
| BD Horizon™ BV510 Mouse IgG1, κ Isotype Control | BD Biosciences | Cat# 562946<br>RRID: AB_2784530 |
| BD Horizon™ BV650 Mouse IgG1, κ Isotype Control | BD Biosciences | Cat# 563231<br>RRID: AB_2869470 |
| BD Horizon™ BB515 Mouse IgG1, κ Isotype Control | BD Biosciences | Cat# 564416<br>RRID: AB_2721017 |
| Donkey anti-Rabbit IgG (H+L) Highly Cross-Adsorbed Secondary Antibody, Alexa Fluor Plus 488 | Invitrogen, Thermo Fisher Scientific | Cat# A21206<br>RRID: AB_2535792 |
| Donkey anti-Mouse IgG (H+L) Highly Cross-Adsorbed Secondary Antibody, Alexa Fluor 594 | Invitrogen, Thermo Fisher Scientific | Cat# A21203<br>RRID: AB_141633 |
| IGF-I Receptor β Antibody | Cell Signaling Technology | Cat# 3027<br>RRID: AB_2122378 |
| Monoclonal Anti-β-actin antibody | Sigma-Aldrich | Cat# A5441<br>RRID: AB_476744 |
| Peroxidase AffiniPure Goat Anti-Rabbit IgG (H+L) | Jackson ImmunoResearch | Cat# 111-035-003<br>RRID: AB_2313567 |
| Peroxidase AffiniPure Goat Anti-Mouse IgG (H+L) | Jackson ImmunoResearch | Cat# 115-035-003<br>RRID: AB_10015289 |
| Phospho-Akt (Ser473) Antibody | Cell Signaling Technology | Cat# 9271<br>RRID: AB_329825 |
| Phospho-IGF-I Receptor β (Tyr1135) (DA7A8) Rabbit mAb | Cell Signaling Technology | Cat# 3918<br>RRID: AB_10548764 |
| Phospho-PLCγ1 (Tyr783) (D6M9S) Rabbit mAb | Cell Signaling Technology | Cat# 14008<br>RRID: AB_2728690 |
| PLCγ1 (D9H10) XP® Rabbit mAb | Cell Signaling Technology | Cat# 5690<br>RRID: AB_10691383 |
| Chemicals, peptides, and recombinant proteins |  |  |
| Albumin from Bovine Serum | Sigma Aldrich | Cat# A8806 |
| BAPTA-AM | Sigma Aldrich | Cat# A1076 |
| BD Pharmingen™ 7-AAD | BD Biosciences | Cat# 559925<br>RRID: AB_2869266 |
| BD Pharmingen™ PE Annexin V | BD Biosciences | Cat# 556422<br>RRID: AB_2869071 |
| BD™ CompBeads Anti-Mouse Ig, κ/Negative Control Compensation Particles Set | BD Biosciences | Cat# 552843 |
| β-Glycerophosphate disodium salt hydrate | Sigma Aldrich | Cat# G9422 |
| BIO5192 | Tocris Bioscience | Cat# 5051/10 |
| CellTrace™ Calcein Red-Orange, AM (Invitrogen) | Thermo Fisher Scientific | Cat# C34851 |
| Collagen, Type I solution from rat tail | Sigma Aldrich | Cat# C3867 |
| D-(+)-Glucose | Sigma Aldrich | Cat# G7021 |
| DAPI dihydrochloride | Himedia | Cat# RM5773 |
| DMEM, High Glucose With 4.5gms Glucose per litre and Sodium pyruvate Without L-Glutamine and Sodium bicarbonate | Himedia | Cat# AT007A |
| EDTA Disodium Salt Dihydrate for molecular biology, 99.5% | Sisco Research Laboratories | Cat# 54448 |
| EGTA | Himedia | Cat# MB130 |
| Fibronectin (Corning®) | BD Biosciences | Cat# 356008 |
| Fluorescein Isothiocyanate–Dextran | Sigma Aldrich | Cat# 46945 |
| Gelatin from bovine skin, Type B | Sigma Aldrich | Cat# G9382 |
| Gibco™ Fetal Bovine Serum, qualified, Brazil | Thermo Fisher Scientific | Cat# 10270106 |
| Histopaque® -1077 | Sigma Aldrich | Cat# 10771 |
| Histopaque® -1083 | Sigma Aldrich | Cat# 10831 |
| Insulin, Human Recombinant (Yeast) | Roche Diagnostics | Cat# 11376497001 |
| Insulin, Human Recombinant | MP Biomedicals | Cat# 0219390080 |
| Iron(III) chloride | Sigma Aldrich | Cat# 157740 |
| Ketamine Hydrochloride 50 mg/ml Injection | Thermis Medicare Ltd., India | N/A |
| L-Glutamine | Sigma Aldrich | Cat# G3126 |
| MG-132 | Sigma Aldrich | Cat# M7449 |
| Okadaic acid | Sigma Aldrich | Cat# O9381 |
| Paraformaldehyde | Himedia | Cat# RM3660 |
| Phenylmethanesulfonyl fluoride | Sigma Aldrich | Cat# P7626 |
| Protease Inhibitor Cocktail | Sigma Aldrich | Cat# P8340 |
| PQ401 | Sigma Aldrich | Cat# P0113 |
| RPMI-1640 With HEPES Buffer and Sodium pyruvate Without Glucose, L-Glutamine and Sodium bicarbonate | Himedia | Cat# AT222A |
| RPMI-1640 With HEPES Buffer and Sodium pyruvate Without L-Glutamine and Sodium bicarbonate | Himedia | Cat# AT157A |
| Sodium Bicarbonate | Sigma Aldrich | Cat# S5761 |
| Sodium Fluoride | Sigma Aldrich | Cat# 106449 |
| Sodium pyrophosphate decahydrate | Sigma Aldrich | Cat# 221368 |
| Streptozotocin | MP Biomedicals | Cat# 100557 |
| U-73122 Hydrate | Sigma Aldrich | Cat# U6756 |
| WESTAR SUPERNOVA ECL substrates for Western Blotting | Cyanagen | Cat# XLS3 0100 |
| Xylazine Hydrochloride | Sigma Aldrich | Cat# X1251 |
| Experimental models: Cell lines |  |  |
| Cell line: Jurkat, Clone E6-1 (Human) | ATCC | Cat# TIB-152™<br>RRID: CVCL_0367 |
| Cell line: 293 [HEK-293] (Human) | ATCC | Cat# CRL-1573™<br>RRID: CVCL_0045 |
| Experimental models: Organisms/strains |  |  |
| Mice: C57BL/6J | NCCS and CSIR-CDRI Animal facility | N/A |
| Software and algorithms |  |  |
| BD FACSDiva™ Software | BD Biosciences | N/A |
| BD CellQuest Pro™ Software | BD Biosciences | N/A |
| FlowJo™ V7.2.5 | BD Biosciences | https://www.flowjo.com/solutions/flowjo/downloads |
| Graphpad Prism | Graphpad | https://www.graphpad.com/scientific-software/prism/ |
| ImageJ | Schneider et al., 2012<br>DOI: 10.1038/nmeth.2089 | https://imagej.nih.gov/ij/download.html |
| Image Lab Software | Bio-Rad | https://www.bio-rad.com/en-in/product/image-lab- |
|  |  | software?ID=KRE6P5E8Z |
| Olympus FV1000 Software | Olympus | N/A |
| Optika Image acquisition Software | Optika | N/A |
| Other |  |  |
| AccuCheck® Blood Glucose Monitoring System | Roche Diagnostics | N/A |
| AccuCheck® Blood Glucose Test Strips | Roche Diagnostics | N/A |
| Amersham™ Hybond® P 0.45µm PVDF blotting membrane | GE Healthcare | Cat# 1060023 |
| BD 1ml Tuberculin Syringe | Becton, Dickinson and Company | Cat# 302100 |
| BD Vacutainer® K2 EDTA Blood Collection Tubes | Becton, Dickinson and Company | Cat# 367863 |
| BD Vacutainer® 3.0 mg sodium fluoride, 6.0 mg Na2EDTA Blood collection tubes | Becton, Dickinson and Company | Cat# 367587 |
| BD Vacutainer® SST™ II Advance Plus Blood Collection Tubes | Becton, Dickinson and Company | Cat# 367956 |
| BD Vacutainer® Flashback Needle for Blood Collection | Becton, Dickinson and Company | Cat# 301747 |
| Dispovan Insulin Syringe (31 G Short needle) | Hindustan Syringes and Medical Devices Ltd. | N/A |
| Oral feeding tube for mice- 20G x 38mm Curved-end | Vishnu Traders, India | N/A |

## RESOURCE AVAILABILITY

### Lead contact

Further information and requests for resources and reagents should be directed to the lead contact, Dr. Madhulika Dixit

### Materials availability

No new resources were generated in this study.

### Data and Code availability

- This study does not report original code.
- The raw data and any additional information required about the data reported in this paper is available from the lead contact upon reasonable request.

## EXPERIMENTAL MODEL AND SUBJECT DETAILS

### Human Cell Lines

HEK293 and Jurkat T-cells were purchased from ATCC. Cultured cells were routinely monitored for appropriate morphology from time to time under a microscope. No new stable, knock-out, knock-in or knock-down cell lines were generated in this study.

### Subject recruitment, inclusion and exclusion criteria

The study was approved by the Institutional Ethics Committee (IEC) of the Indian Institute of Technology Madras (IIT Madras) in accordance with the ‘National Ethical Guidelines for Biomedical and Health Research involving human participants’ of the Indian Council of Medical Research (ICMR), Government of India. Recruitment of the participants and sample collection took place in the Out-Patient facility of the Institute hospital of IIT Madras, Chennai. A total of 20 drug naïve healthy Asian Indian men within the age range of 20-45 years with no prior history of diabetes, were screened through a standard 75g oral glucose tolerance test (OGTT), following written informed consent. Normal glucose tolerance (NGT) individuals with fasting plasma glucose ≤ 99 mg/dl and 2-hour post load plasma glucose of ≤ 139mg/dl were recruited in the study. Individuals who were prediabetic (fasting plasma glucose between 100-125 mg/dl and/or 2-hour post load plasma glucose of 140-199 mg/dl) and those who were overtly diabetic (fasting plasma glucose ≥ 126 mg/dl and/or 2-hour post load plasma glucose ≥ 200 mg/dl) were excluded from the study. Other exclusion criteria that were applied are as follows: hypertension (>139/90 mmHg, SBP/DBP), established cardiovascular disorders, previously diagnosed diabetes, cancer, any form of medication, smoking, chronic alcoholism, surgery in the past 1 year and recent infection. A total of sixteen NGT subjects were recruited in this study. Biochemical parameters such as plasma glucose (Hexokinase method), serum insulin (Electrochemiluminescence Immuno Assay i.e. ECLIA), serum lipid profile (Total Cholesterol by CHOD-PAP assay, Triglycerides by GPO-PAP assay, HDL by direct immuno-inhibition method, LDL and VLDL by calculation), as well as total and differential WBC counts (Impedance based differential counting method) were measured. Anthropometric parameters such as height and weight were measured for each individual. Blood pressure was measured using a semi-automated sphygmomanometer from the non-dominant arm of the individuals in a relaxed state prior to collecting the fasting blood sample.

## METHOD DETAILS

### Peripheral Blood Mononuclear Cell (PBMC) isolation for functional assays

Peripheral blood mono-nuclear cells (PBMCs) were isolated by density gradient centrifugation at room temperature by layering blood over equal volume of Histopaque-1077. Buffy coat was collected and washed once with 1X phosphate buffered saline (PBS) [137mM NaCl, 2.7mM KCl, 10mM Na_2_HPO_4_, 1.8mM KH_2_PO_4_, pH 7.4] and PBMCs were pelleted by centrifugation at 1800 rpm for 10 minutes. PBMC pellet was subjected to RBC lysis in 1ml of 1X RBC lysis buffer [150mM NH_4_Cl, 10mM NaHCO_3_, 100nM EDTA, pH 7.4] for 10 minutes at room temperature, followed by another wash with 1XPBS.

### Cell culture

Freshly isolated PBMCs were used for cell adhesion and trans-well migration assays. For experiments involving *ex-vivo* treatment of fasting PBMCs with insulin, agonist treatment with varying concentrations of endotoxin-free insulin was performed for desired durations in RPMI-1640 basal medium lacking fetal bovine serum (FBS), but supplemented with 5mM glucose and 0.1% bovine serum albumin (BSA) (referred to as RPMI basal medium henceforth). Jurkat T-cells were maintained in RPMI-1640 growth medium supplemented with 10 mM glucose and 10% FBS. They were cultured at 37°C in presence of 5% CO_2_ for a maximum of 6 passages after revival. Jurkat cells were serum starved for 6 hours prior to experimental treatment with insulin. Serum starvation and insulin treatment of Jurkat cells was performed in RPMI-1640 basal medium.

### Adhesion assay

Wells of 48-well cell culture dishes were coated with fibronectin (25 μg/mL), collagen (0.01% w/v) or gelatin (0.2% w/v) for 1 hour at 37°C prior to the start of adhesion experiments. Freshly isolated PBMCs or Jurkat cells (1X 10^5^ cells) were seeded onto the coated wells, in RPMI-1640 basal medium in the absence of FBS or BSA and were allowed to adhere for 3 hours in cell culture compatible CO_2_ incubator at 37°C. Non-adherent cells were removed through washing of culture wells with 1X PBS twice. Adherent cells were fixed with 4% paraformaldehyde in 1X PBS (pre-warmed to 37°C) at room temperature for 5 minutes, followed by washing with 1X PBS twice before staining them with 1μg/mL DAPI for 2 minutes at room temperature. Images were taken using an Olympus IX51 fluorescence microscope. Multiple fields of view per well were analysed. Data are represented as average number of cells adhered per field of view for a minimum of four independent experiments. The acquired images were analyzed using ImageJ software.

### Trans-well migration assay

Trans-well migration filters of 8 μm pore size were coated with fibronectin (25 μg/mL) on both sides at 37°C for 1 hour. This was followed by seeding of PBMCs (2 × 10^5^ cells) onto the upper chamber of the trans-well filter in RPMI-1640 basal medium. Cells were incubated for 16 hours at 37°C in 5% CO_2_. Following incubation, the cells on the upper side of the membrane were carefully removed using a sterile cotton swab, and the cells that migrated to the lower side of the membrane were fixed with 4% paraformaldehyde and stained with DAPI (1μg/mL) for image acquisition. The lower side of the membrane was imaged in multiple fields of view. Data are represented as the average number of cells migrated per field of view. The acquired images were analyzed using ImageJ software.

### Flow-cytometry

In all flow-cytometry experiments, acquisition was performed on BD CANTO II/ARIA III/Calibur. A minimum of 1X10^5^ events were acquired in all experiments. Amount of primary and secondary antibodies used for staining were according to the manufacturer’s instructions. Single stain compensation controls were used during multi-colour acquisitions. Appropriate isotype controls were used wherever necessary. In secondary staining experiments, a negative control with only secondary antibody staining was performed to confirm specificity. Data was analysed using FACS DIVA or Flow-Jo software and data is represented as percentage of cells positive for a given protein within the gated population. Analyses involving PBMCs were performed by gating for the lymphocyte-monocyte fraction. Average expression of a given protein in each cell is represented as the geometric mean of the fluorescence intensity (MFI) of the population positive for that specific protein.

### Surface staining of PBMC and Jurkat cells

Blocking was performed with 0.1% BSA and 5% FBS per sample in 1X PBS for 30 minutes at room temperature. Respective antibodies were added and incubated at 4°C or at room temperature for 30 minutes, as per the manufacturer’s instructions. After staining, the cells were washed twice with 1X PBS and fixed with 0.1% paraformaldehyde in 1X PBS and stored at 4°C until acquisition.

### Surface staining of Jurkat cells for activation specific integrin antibodies

Following serum starvation, Jurkat cells were incubated with 10nM insulin in RPMI-1640 basal medium containing 3mM EDTA. Cells were fixed by adding equal volume of pre-warmed 4% paraformaldehyde in 1X PBS to the cell suspension for 10 minutes at room temperature at the end of insulin treatment. Blocking was performed with 0.5% BSA for 30 minutes at room temperature. Cells were then incubated with fluorochrome conjugated primary antibodies, Ts2/16 β1 and m24 β2 integrin clones specific for activated integrins, for an hour at 4°C in 1X PBS. Following washing with 1X PBS, cells were fixed with pre-warmed 0.1% paraformaldehyde in 1X PBS for 10 minutes at room temperature prior to acquisition.

### Intracellular staining for pIGF-1R in Jurkat cells

Cells were serum starved for 6 hours and treated with 1µM proteasome inhibitor MG132 for 4 hours (during serum starvation) at 37°C prior to insulin treatment. Cells were fixed by adding equal volume of pre-warmed 4% paraformaldehyde in 1X PBS to the cell suspension for 10 minutes at room temperature at the end of insulin treatment. Fixed cells were pelleted by centrifugation at 1800 rpm for 10 minutes and were permeabilised by re-suspending them in 90% ice cold methanol for 10 minutes on ice. Permeabilization was stopped with RPMI-1640 medium supplemented with 10% FBS and the cells were washed with 1XPBS and blocked with 0.1% BSA for 30 minutes at room temperature. Primary antibody for Tyr^1135^ Phospho-IGF-1R was then added to the blocking suspension and incubated for 30 minutes at room temperature. Cells were washed with 1X PBS at room temperature and re-suspended in 1X PBS followed by incubation with 2µg Alexa fluor 488 conjugated secondary antibody for 30 minutes at 4°C. Finally, the cells were washed with 1X PBS and were fixed with pre-warmed 0.1% paraformaldehyde before acquisition.

### Cell viability assay

PBMCs were suspended in 100μl of binding buffer [10mM HEPES, 140 mM NaCl, 2.5mM CaCl_2_, pH 7.4] and stained with 7-AAD and PE conjugated Annexin V as per manufacturer’s instruction for 20 minutes at room temperature. After incubation, the suspension was diluted to 500µl with 1X HEPES buffer and acquired via flow cytometry within 30 minutes.

### Inhibitor treatments

At the end of 6 hours of serum starvation, prior to insulin treatment, Jurkat cells were pre-treated with the following inhibitors in relevant experiments for required duration as listed below:

a. 20µM PQ401, an irreversible IGF-1 receptor (IGF-1R) auto-phosphorylation inhibitor, for 45 minutes.
b. 1µM U73122, a reversible PLC inhibitor, for 1 hour.
c. 5µM BIO5192, a potent inhibitor of α4β1 integrin for 1 hour.
d. 10µM BAPTA-AM, an intra-cellular calcium chelator, for 45 minutes.
e. To determine the effect of extra-cellular cations, Jurkat cells were treated with insulin, following which, the adhesion assay on fibronectin was performed either in the presence or absence of 3mM EDTA.

Corresponding solvents of the inhibitors were used as vehicle controls.

### Western blotting

Jurkat cells were serum starved for 6 hours in RPMI basal medium with 0.1% BSA. For the PLC-γ1 and Akt phosphorylation time-course experiments, following serum starvation, 10nM insulin treatment was given for respective time duration. For the PLC-γ1 phosphorylation experiments following IGF-1R phosphorylation inhibition, the cells were treated with 20 µM PQ401 for 45 minutes at the end of serum starvation, before insulin treatment. At the end of the insulin treatment in both experiments, the cells were harvested in medium containing 1mM sodium orthovanadate, pelleted and snap frozen in liquid nitrogen. For the IGF-1R phosphorylation experiments, cells were treated with 1µM proteasome inhibitor MG132 for 4 hours (during serum starvation), prior to insulin treatment. Immediately after insulin treatment, cells were fixed by adding equal volume of pre-warmed 4% paraformaldehyde in 1X PBS containing 1mM sodium orthovanadate to the cell suspension for 10 minutes at room temperature, pelleted and frozen at −80°C.

Cells were lysed in Laemmli buffer supplemented with 5 mM EDTA, 1 mM EGTA, 5 mM sodium orthovanadate, 50 mM NaF, 10 mM β-glycerophosphate, 10 mM sodium pyrophosphate, 10 nM okadaic acid, 5mM PMSF, and 1X protease and phosphatase inhibitor cocktails. Lysates were resolved by SDS PAGE and transferred to PVDF membrane, followed by blocking with 5% BSA in TBST [137 mM NaCl, 2.7 mM KCl, 19 mM TRIS base (TBS) and 0.1% tween-20]. Primary and secondary antibody treatments for phospho-proteins and total proteins were carried out according to manufacturer’s instruction. The blots were developed using enhanced chemiluminescence using Cynagen Westar Supernova detection kit and imaged using ChemiDoc^TM^ system from Bio-Rad. The blots were then stripped and re-probed for total proteins. Briefly, stripping buffer [62.5mM Tris-HCl, 2% SDS and 100mM β-mercaptoethanol, pH 6.7] was pre-warmed to 65°C and added to the blots which were incubated in the buffer with agitation for 45 minutes at 65°C. Following stripping, the blots were washed twice with 1X TBS and twice with 1X TBST for 10 minutes in each wash. Then the blots were blocked again and probed for total proteins, developed and imaged as described above. The images were analysed using ImageJ software.

### Animal Experiments

The study was approved by the Institute Animal Ethics Committee at the National Centre for Cell Science [IAEC/2018/B-268(I)], Pune, India and in Central Drug Research Institute [IAEC/2014/43], Lucknow, India, in accordance with the Committee for the Purpose of Control and Supervision of Experiments on Animals (CPCSEA) guidelines, Government of India. Male C57BL/6 mice within 8-12 weeks of age were maintained on standard pelleted rodent diet and filtered water *ad libitum* in 12-hour light and dark conditions until experimentation.

### OGTT, PBMC isolation and adhesion assay

Mice were fasted for 6 hours, with access only to water, following which, fasting blood sample was collected by retro-orbital bleeding. Glucose solution equivalent to 2gm/kg body weight of the animal was fed through oral gavage and blood sample was again collected 1-hour post glucose load. 100µl blood sample was diluted to 1ml with 1X PBS and buffy coat was isolated from the diluted blood sample by layering over 1ml Histopaque-1083 followed by density gradient centrifugation for 30 minutes at room temperature. Isolated PBMCs were washed twice with 1X PBS and were subsequently pelleted by centrifugation at 1800 rpm for 10 minutes. Isolated PBMCs (1×10^5^) were seeded onto wells coated with fibronectin (25µg/ml), in RPMI-1640 basal medium in the absence of serum and were allowed to adhere for 3 hours. Non-adherent cells were washed with 1X PBS twice. Adherent cells were fixed with pre-warmed 4% paraformaldehyde in 1X PBS for 5 minutes at room temperature, washed twice with 1X PBS and then stained with 1μg/mL DAPI in 1X PBS for 2 minutes at room temperature. Multiple field views were imaged per well. Data are represented as average number of cells adhered per field of view. The images were analyzed using ImageJ software.

### Single dose streptozotocin (STZ) injection and *ex-vivo* insulin treatment of PBMCs

Healthy C57BL/6 male mice were fasted overnight with access to water, following which they were weighed and 200 mg/kg body weight streptozotocin (STZ) freshly prepared in citrate buffer [25mM citric acid and 25mM trisodium citrate dihydrate solution, pH 4.5] was injected intra-peritoneally. For control animals, only buffer was injected. STZ injected animals were maintained on standard pelleted rodent diet and filtered water *ad libitum* for 72 hours, at the end of which, their random blood glucose levels were determined using tail clipping with a glucometer. In the STZ injected group, only animals with random blood glucose levels greater than 300 mg/dl, due to loss of insulin, were used for the experiments. OGTT, PBMC isolation and adhesion assays were performed as described above. For the *ex-vivo* insulin treatment experiments, fasting PBMCs from the animals were incubated with 10nM insulin for 1-hour in RPMI-1640 basal medium, following which, adhesion assay on fibronectin was performed.

### Intra-vital Microscopy

Experimental animals were procured from the National Laboratory Animal Centre of CSIR-CDRI Lucknow. Donor C57BL/6J mice were fasted overnight (10-12 hours) and were administered 2 gm/kg D-Glucose (Dextrose), through oral gavage. Blood glucose was monitored before and after 1-hour of glucose administration using Accu-Chek glucometer (Roche Diagnostics, India). Fasting and post glucose load PBMCs were isolated from freshly obtained whole blood from the donor mice using density gradient separation with Histopaque-1083. Blood was collected by cardiac puncture from donor mice in EDTA-coated centrifuge tubes. Anticoagulated diluted blood was layered onto two ml of pre-warmed Histopaque for isolation of PBMCs through density gradient centrifugation. PBMCs were collected and re-suspended in PBS followed by labelling with Calcein red orange (2.5µg/ml). Recipient C57BL/6J male mice were anesthetized with ketamine (80 mg/kg) and xylazine (20 mg/kg). FITC-labelled Dextran (2mg/mice) and Calcein red orange labelled donor PBMCs (1×10^9^/kg) were injected through retro-orbital plexus to visualize the blood vessels and PBMCs respectively. Mesenteric artery of the recipient mice was exposed through an abdominal wall incision and a Whatmann filter paper (2mm x 0.5mm) saturated with 5% FeCl_3_ (w/v) solution was applied topically for 3 minutes for inducing the endothelial/vessel injury. Adherence of labelled donor PBMCs to the exposed basement membrane of the mesenteric artery of the recipient mice was monitored under intra-vital microscope [Olympus FV1200 (BX61Wi), Japan] coupled to a CCD camera [28, 29]. The images were captured and quantified using FV1000 (Olympus) and image J software.

### Statistical analyses

Statistical analyses were performed using Graphpad Prism software. All data is represented as mean ± SEM. Determination of normal distribution was done through Kolmogorov-Smirnov (KS) test. Paired and unpaired two-tailed Student’s t-test were applied wherever applicable. One-way ANOVA followed by Dunnett’s post-hoc test was applied while comparing multiple conditions to a single control condition and Tukey’s post-hoc test was applied for comparison of multiple conditions with every other condition. Statistical significance was achieved for p values less than 0.05. Specific statistical tests applied for each of the experiments are described in the respective figure legends.

