## Supplementary figures and images for "Oral glucose feeding enhances adherence of quiescent lymphocytes to fibronectin via non-canonical insulin signalling"

### Supplemental Figure 1

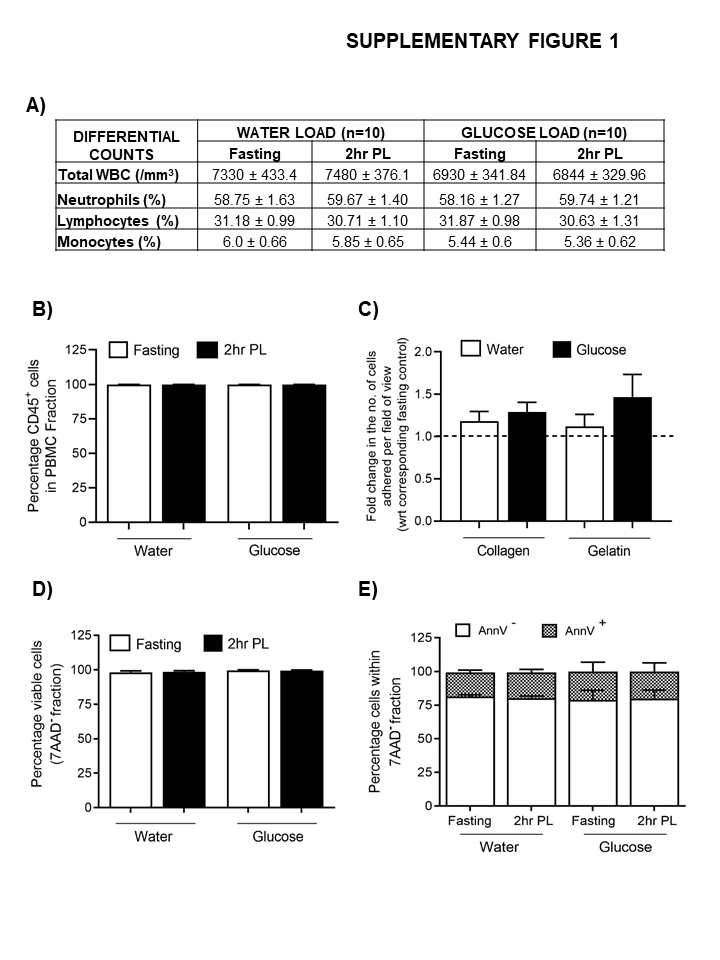

### Supplemental Figure 2

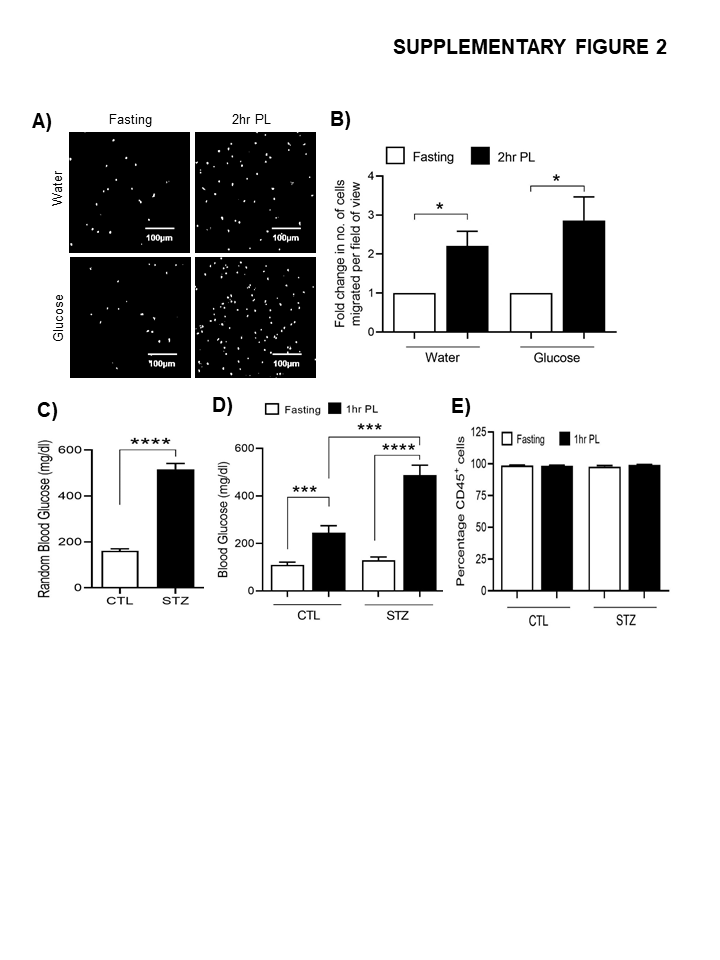

### Supplemental Figure 3

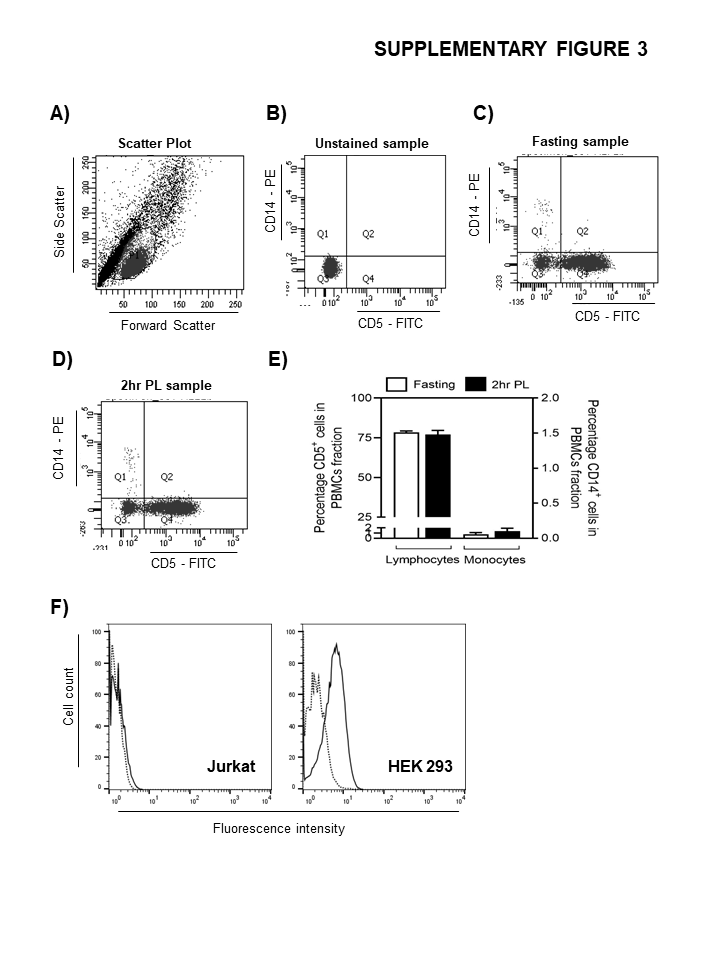

### Supplemental Figure 4

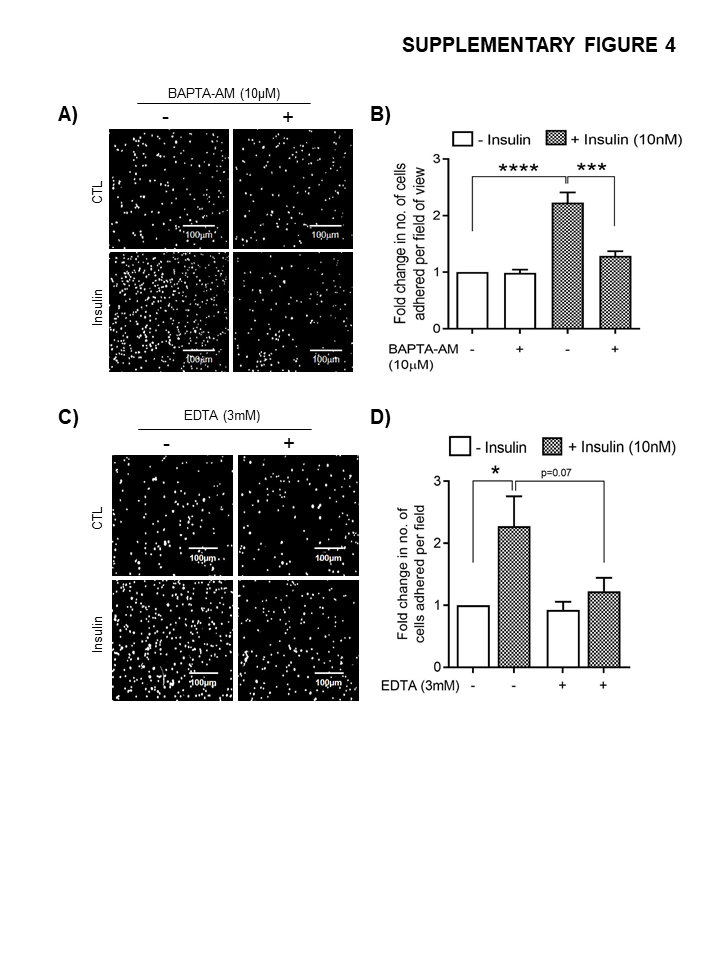

### Supplemental Figure 5

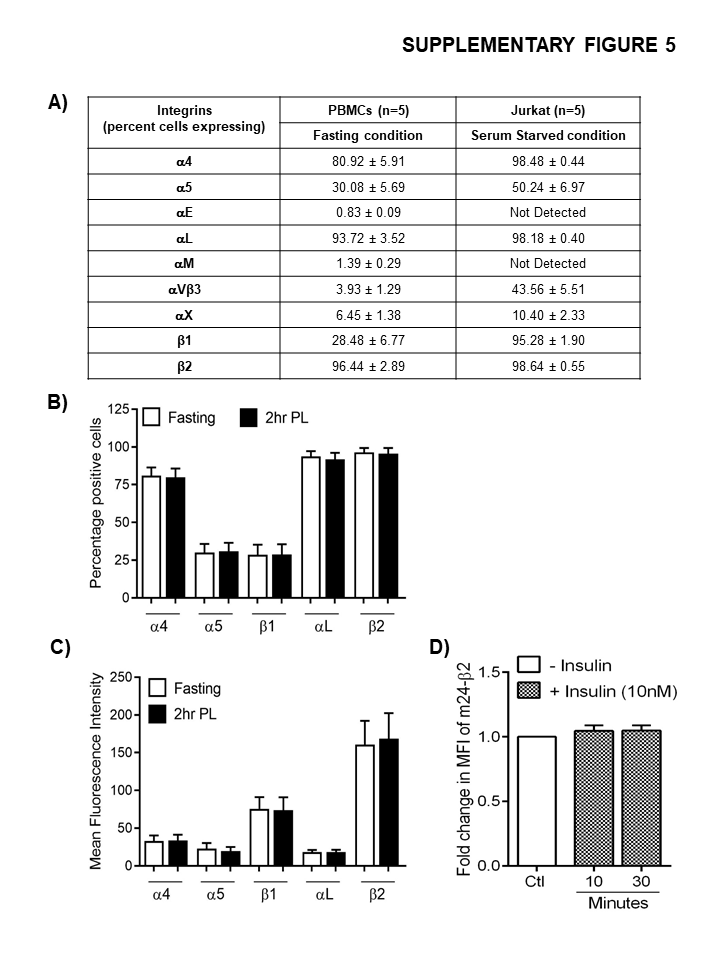
